# Gene regulatory networks define human airway epithelial cell types and their distinct responses to type I interferon

**DOI:** 10.64898/2026.05.09.724010

**Authors:** Anthony T. Bejjani, Alec P. Pankow, Ava R. Lamberty, Jorg J.A. Calis, Kira A. Griswold, Ethan Iverson, Andy Kuo, Akshata N. Rudrapatna, Skyler A. Uhl, Roosheel S. Patel, Oded Danziger, Philip Cohen, Erika A. Barrall, Emma J. DeGrace, Fady Gorgy, Ngoc Pham, Zi F. Yang, Joseph A. Wayman, Alexander Katko, Allison Boboltz, Gregg A. Duncan, William J. Zacharias, Ivan Marazzi, Matthew T. Weirauch, Leah C. Kottyan, Christopher Benner, Margaret A. Scull, Brad R. Rosenberg, Emily R. Miraldi

## Abstract

The human airway epithelium (**HAE**) is composed of diverse cell types that coordinate essential functions and host defenses. Among these defenses, interferons (**IFNs**) are central to antiviral programs. However, the gene regulatory networks (**GRNs**) governing HAE cellular identities and their IFN responses are incompletely defined. At single-cell resolution, we characterized the transcriptomes and accessible chromatin landscapes of HAE cell types, at steady state and following IFNβ stimulation. The resulting scRNA-seq and snATAC-seq data informed genome-scale GRN construction and inference of transcriptional circuits underlying cell identities. In response to IFN, we identified a shared transcriptional program across HAE cell types, and an expanded set of interferon-responsive genes exhibiting cell type-associated expression patterns. Cell type-associated transcription factors and chromatin accessibility contribute to distinct IFN-responsive gene expression programs. Together, these data provide a blueprint for molecular regulation of complex HAE responses and a foundation for therapeutic strategies to enhance host antiviral defense.

## Introduction

Respiratory viral infections remain a major global health challenge, substantially contributing to morbidity and mortality across all age groups. Seasonal epidemics, along with emergent pathogens such as SARS-CoV-2, cause millions of severe infections annually, often leading to pneumonia, exacerbation of chronic lung disease and long-term pulmonary impairment^1^. Further, effective antiviral therapeutics for respiratory infections are limited and largely pathogen specific^2^.

The human airway epithelium (**HAE**) is the primary target for infection by respiratory viruses. As such, the HAE has evolved myriad defense mechanisms to protect against inhaled pathogens and promote their clearance. Multiple cell types comprise the HAE, and these cell types have distinct distributions throughout airways as well as unique roles in both airway homeostasis and innate defense. For example, goblet cells produce mucus that traps debris, ciliated cells propel mucus towards the oropharynx for clearance, and basal cells continuously renew the epithelium, ensuring its integrity. Single-cell transcriptomic (**scRNA-seq**) studies have refined our understanding of HAE composition^3–5^, enabling the identification of ionocytes and deuterosomal cells, which play important roles in health and disease, including cystic fibrosis, respiratory infections and chronic obstructive pulmonary disease^7–11^. Still, the precise molecular programs governing diverse HAE component cell types, particularly during innate antiviral responses, remain incompletely understood.

Type I and type III interferons (**IFNs**) are key contributors to HAE innate antiviral defense during respiratory virus infection. Defects in IFN signaling lead to severe outcomes following respiratory virus infection, for example, in individuals with inborn errors in pathogen sensing or IFN signaling pathways^12–18^ or autoantibodies targeting type I IFNs (**IFN-I**)^17,19,20^. Similarly, animals lacking IFN receptor components or downstream IFN signaling factors are exceedingly susceptible to viral infections, often exhibiting elevated viral titers, disseminated infection and increased mortality^21^. In agreement, IFN gene signatures are prominently detected from scRNA-seq analyses of HAE exposed to pathogenic insults^22–25^.

Given the critical host defense roles of IFNs, it is not surprising that IFN-I and IFN-III have been evaluated as potential therapeutics for viral infections. Although pegylated IFN-α2a is moderately efficacious in the treatment of hepatitis B and C virus infections, administration is accompanied by severe side effects^26–29^. IFN-I and IFN-III have also been evaluated as potential therapeutics for respiratory virus infections, although challenges associated with mucosal delivery, increased susceptibility to bacterial infections, and potential negative impacts on tissue recovery have curtailed their widespread implementation^30,31^. An improved understanding of IFN responses in HAE may enable novel therapeutic strategies that incorporate potent antiviral features and minimize side effects and inflammatory damage.

The type I interferons signal through the interferon alpha/beta receptor (**IFNAR**) via the JAK-STAT pathway to enact antiviral and immunomodulatory programs^32–35^. Upon cytokine binding, receptor-associated tyrosine kinase 2 (TYK2) and Janus kinases (**JAKs**) become phosphorylated and in turn phosphorylate signal transducers and activators of transcription (**STATs**) STAT1 and STAT2. Phosphorylated STAT1 and STAT2 preferentially form heterodimers and associate with interferon regulatory factor 9 (**IRF9**) to form the interferon stimulated gene factor 3 (**ISGF3**) complex. The ISGF3 complex translocates to the nucleus, where it binds to interferon-stimulated response elements (**ISREs**) and drives transcription of interferon-stimulated genes (**ISGs**). Additionally, phosphorylated STAT1 can form homodimers (**STAT1:STAT1**) and bind to gamma interferon activation site (**GAS)** elements to induce gene expression^36^. IFN-induced gene expression programs establish an antiviral state accompanied by additional antiproliferative, proapoptotic and immune crosstalk effects. Some ISG products act as direct antiviral effectors that restrict viral life cycle processes via diverse mechanisms. Other ISG products include cytokines, chemokines and antigen presentation pathway components, which promote a multifaceted host response by recruiting additional immune cells to infection sites, thereby linking innate and adaptive immune activities^37–39^.

Here, we leverage a well-differentiated HAE tissue model at the air-liquid interface to precisely characterize the early dynamics of the HAE response to Type I interferon IFNβ and infer the gene regulatory networks (**GRNs**) underlying this important process. GRNs describe the control of gene expression by TFs. Given the heterogeneity of cell types composing the HAE, single-cell data is essential for GRN accuracy and its downstream application to identify new targets, modulate epithelial defenses and improve outcomes in respiratory viral disease. We therefore constructed genome-scale GRNs for HAE from single-cell-resolved transcriptomics and chromatin accessibility measurements (scRNA-seq, snATAC-seq and sn-multiome-seq) from an IFNβ time course (**Fig. 1A**). Context-specific TF binding predictions from chromatin accessibility data improve GRN inference from gene expression^40–43^. In this study, we simulated *maxATAC* deep neural network models^44^ to generate genome-wide, cell-type and timepoint-resolved “*in silico* TF ChIP-seq” data to guide GRN inference. Although individual regulators have previously been associated with HAE cell types (e.g., FOXN4 in deuterosomal cells^45^), our GRN predicts nearly one hundred TF regulators of HAE cell-type identities and connects them to target genes and cell-type-specific functions. In response to IFNβ, the GRN uncovered both expected (e.g., ISGF3) and novel, cell type-associated regulators. The GRNs and underlying data provide functional and regulatory insights into the diverse cell types composing the HAE and their distinct responses to interferon.

**Figure 1:**
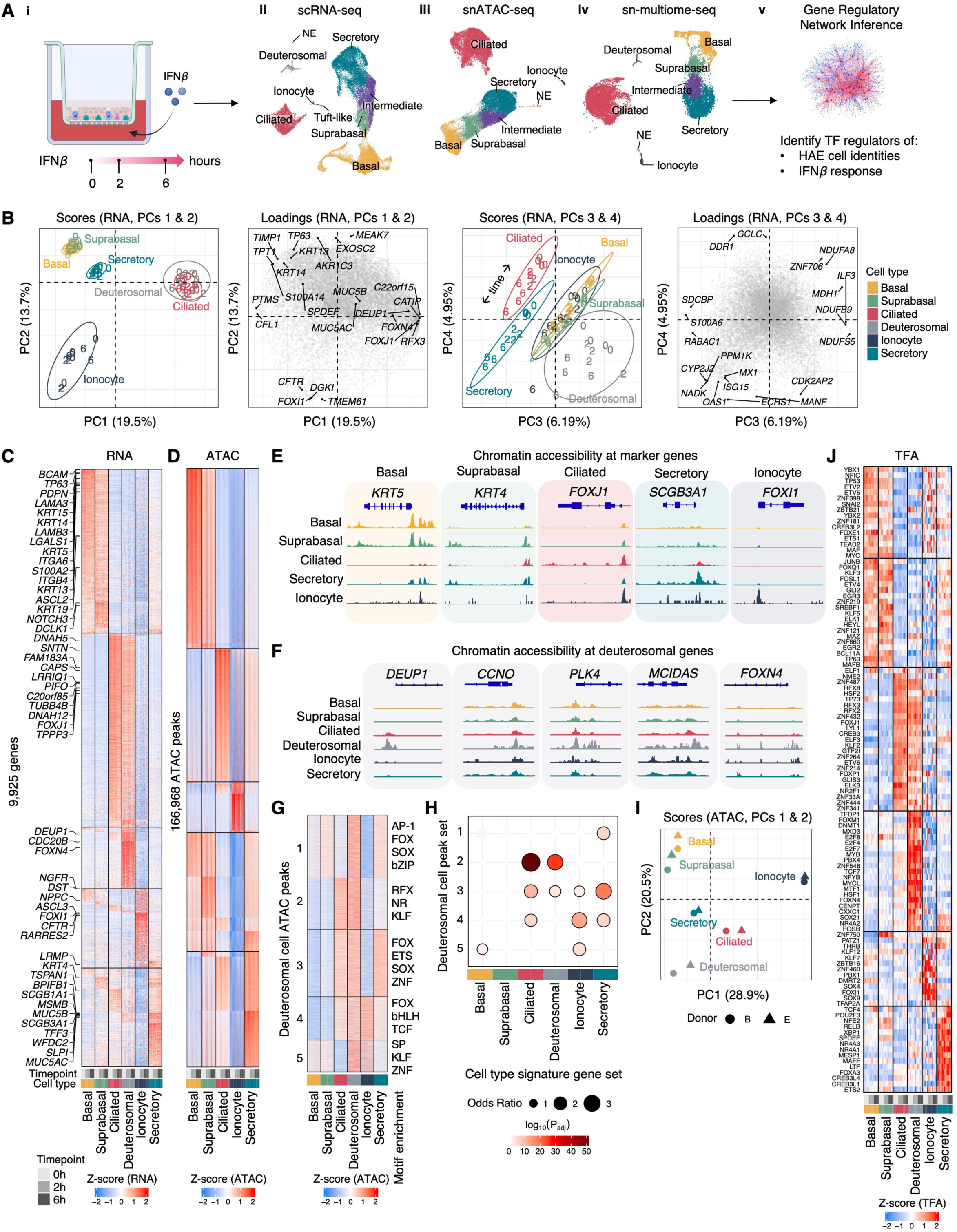
Single-cell resolution transcriptome and accessible chromatin landscapes of HAE component cell types. (**A**) (**i**-**v**) Study design, (**ii**-**iv**) UMAP visualization of (**ii**) scRNA-seq, (**iii**) snATAC-seq and (**iv**) sn-multiome-seq cell type annotations. (**B**) Principal component analysis (**PCA**) of pseudobulk (cell type, time point, donor) gene expression data. In the scores plots, pseudobulk samples are colored by cell type, with timepoint indicated (“0”, “2”, or “6” hours post-IFN) for each of four donors. Ellipses represent 95% confidence intervals for each cell type. Loadings plots depict gene contributions to the PCs, with select genes labeled. Axis labels include percent variance explained by each PC in parentheses. (**C**) Normalized pseudobulk gene expression (RNA, z-scaled) of 9,925 signature genes (P_adj_ < 0.1, |log_2_(FC)| > 0.58, **Methods**) distinguishing the major epithelial cell types; labels indicate cell type marker genes described in Hewitt and Lloyd^46^ (4 donors per cell type and 3 timepoints). (**D**) Normalized pseudobulk (cell type, time point, donor) chromatin accessibility (z-scaled) of 166,968 differential peaks (|log_2_(FC)| > 0.58) distinguishing five HAE cell types (3 donors per cell type and 3 timepoints). (**E**) Chromatin accessibility (from snATAC-seq assay) at select cell type marker gene loci across HAE cell types. Box colors indicate the associated cell type for each marker. Coverage signal normalized per million snATAC-seq fragments. (**F**) Chromatin accessibility (from sn-multiome-seq) of select deuterosomal cell marker genes across the major epithelial cell populations. Coverage signal normalized per million snATAC-seq fragments. (**G**) Normalized pseudobulk (cell type, time point, donor) chromatin accessibility of deuterosomal cell peaks (from sn-multiome-seq, n = 2 donors) across HAE cell types. Each cluster is annotated (right-hand-side) by TF families whose motifs are significantly enriched in that cluster (FDR = 5%, Fisher’s exact test). (**H**) Enrichment of accessible chromatin regions detected in deuterosomal cells (from **G**) proximal to nominally detected genes (±2kb of TSS) in cell type signature gene sets (from **C**) (P_adj_ < 0.05 indicated by black circle outline, Fisher’s exact test). (**I**) PCA of pseudobulk ATAC-seq signal from sn-multiome-seq; pseudobulks were down-sampled to eliminate library-size (technical) variability. (**J**) GRN-estimated protein TF activities (**TFAs**) of “core” TFs predicted to control cell type-specific gene signatures at steady-state. HAE schematics in (**A**) generated with BioRender.

## Results

### Single-cell analyses define transcriptional and chromatin landscapes for the component cell types of the human airway epithelium

We used single-cell technologies to characterize gene expression and chromatin accessibility of HAE component cell types, at steady-state (0 hours) and in response to basolateral IFNβ stimulation (2 and 6h post-treatment) in HAE cultures generated from four donors (**Fig. 1A**). Across the time course, we recovered high-quality data for 30,786 cells from scRNA-seq, 45,561 nuclei from snATAC-seq, and 13,522 nuclei from sn-multiome-seq. Based on canonical marker gene expression^3,6,10,46–48^, we annotated component HAE cell types: basal, suprabasal, ciliated, secretory (including both putative club and mucous/goblet cells with overlapping gene signatures^46^), ionocyte, tuft-like, neuroendocrine (**NE**) and deuterosomal cells; “intermediate” cells exhibited mixed basal and secretory gene expression programs, consistent with previously described transition differentiation states^49,50^ (**Fig. 1A**, **Fig. S1A-C**).

We next assessed sources of gene expression variation, aggregating scRNA-seq data into time point- and donor-resolved pseudobulk profiles for each of the six most abundant HAE cell types (**Table S1**). In principal component analysis (**PCA**) of gene expression data, cell type was the greatest source of variation, dominating principal components (**PCs**) 1-3, while PC4 captured dynamic gene expression responses to IFNβ (**Fig. 1B**). Many expected genes contributed to the PC loadings. For example, *FOXI1* and *CFTR* contributed to the ionocyte-defining PC2, while canonical ISGs (e.g., *ISG20*, *OAS1*, *TNFSF10*) composed the IFNβ timepoint-dependent PC4. Differential gene expression analysis across cell types identified 9,925 genes distinguishing basal, suprabasal, ciliated, deuterosomal, ionocytes and secretory cells at steady- state (**Fig. 1C**).

Applying scRNA-seq cell type annotations to the snATAC-seq data via label transfer^51^, we defined accessible chromatin profiles for all major HAE cell types except deuterosomal cells (**Fig. 1A**, panel iii, **S1A**). Across the five HAE cell types detected, we identified 453,200 accessible chromatin regions (“peaks”), of which 166,968 (37%) were differentially accessible between cell types at steady state (**Fig. 1D**, **Table S2A**). Cell type annotations were supported by chromatin accessibility proximal to cell-type marker gene loci (e.g., elevated accessibility at *KRT5* in basal cell types, *FOXJ1* in ciliated cells, *SCGB3A1* in secretory cells and *FOXI1* in ionocytes, **Fig. 1E**). Mirroring the transcriptome data, cell type was again the greatest source of variation in the chromatin accessibility data, and grouping of biological replicates supported data reproducibility (**Fig. S1D**).

Despite their apparent underrepresentation in snATAC-seq data, deuterosomal cells constituted a similar fraction of total cells in the scRNA-seq and sn-multiome-seq datasets (**Fig. 1A**). Deuterosomal cells are specialized progenitors for multiciliated cells^52^ that repurpose components of the G2/M machinery to drive the extensive centriole amplification required for assembling multiple motile cilia^53–55^. Consistent with their utilization of cell cycle components for cilia assembly, most (∼75% of either scRNA or sn-multiome-derived populations) deuterosomal cells were classified as “G2M”, in gene expression cell cycle phase predictions (**Fig. S1E**). Deuterosomal marker gene transcripts were similarly detected across scRNA and sn-multiome-seq technologies (**Fig. S1F**).

To elucidate the chromatin state of deuterosomal cells, we evaluated chromatin accessibility in the 114 deuterosomal cells detected in the sn-multiome-seq data. We observed elevated chromatin accessibility proximal to deuterosomal markers and genes previously implicated in the developmental trajectory from deuterosomal to ciliated cells^56^ (**Fig. 1F**). In total, we detected 11,745 peaks in deuterosomal cells, which were reproducibly detected across the sn-multiome-seq donors and clustered into five patterns of accessibility across the six HAE cell types (**Fig. 1G**, **Table S2B**). The “cluster 1” peaks were most exclusively accessible in deuterosomal cells, while the remaining clusters were shared with ciliated, ciliated and secretory, all differentiated HAE cell types or ionocytes. The deuterosomal cell peak clusters varied in their TF motif enrichments, suggesting unique and shared gene regulatory mechanisms. The peak clusters also varied in their proximity to the genes defining each HAE cell types at steady state, which we refer to as “signature genes” (**Fig. 1H**, refer to “signature genes” definition **Methods**). For example, “cluster 2” peaks, which have the highest accessibility in deuterosomal and ciliated cells, were proximal to deuterosomal and ciliated cell signature genes and enriched for RFX family TF motifs, consistent with the known role of RFX family members in ciliated and deuterosomal cell regulation^57,58^. Global analysis of the multiome-seq-derived chromatin accessibility data provided a complementary view: the accessible chromatin profiles of deuterosomal cells were most similar to, but also distinct from, ciliated and secretory cells, as biological replicates grouped separately for each of the cell types (**Fig. 1I**).

We next used the *Inferelator*^40,59,131^ to construct genome-scale GRNs for the HAE cell types. The *Inferelator* models gene expression as a function of TF activities (**TFAs**), using TF binding site (**TFBS**) predictions, derived from snATAC-seq, to approximate protein TFAs and guide gene expression modeling. Both cell type differences and IFN response dynamics provided useful gene expression variation for GRN inference (**Table S3A**). For rare populations, we aggregated sc-transcriptomes from biological donors and/or timepoints to achieve necessary minimum pseudobulk library sizes. In contrast, for abundant secretory, basal and intermediate populations we used high-resolution clustering to leverage additional gene expression heterogeneity for GRN inference (**Fig. S1B**, **S2A-B**, **Table S1B**). In total, the GRN describes 123,993 regulatory interactions between 722 TFs and 12,417 genes, a model complexity (i.e., size) determined by predictive performance (**Fig. S2C**). Using the GRN, we recalculated TFAs, based on the expression of the TFs’ predicted gene targets (as opposed to the TFs’ mRNA expression)^59,60^ (**Table S3B**). Across the six cell types at steady state, we detected 109 TFs with differential TFA, including 17 TFs uniquely defining deuterosomal cells (**Fig. 1J**).

### Gene regulatory networks define HAE cell types at steady-state

We first explored how TFs orchestrate HAE cell type identity at steady state. For each of the six most abundant HAE cell types, we derived “core” GRNs^61^, defined here as a set of TFs and their regulatory interactions, predicted to control signature genes and cell type identities at the 0h timepoint:

#### Deuterosomal and ciliated cells

Recently identified in scRNA-seq studies^3–6^, deuterosomal cells transition to terminally differentiated ciliated cells through a sequential process involving the biogenesis, amplification, apical migration and docking of basal bodies, and cilium assembly^52,62–64^. This process is reflected in deuterosomal and ciliated cell transcriptomes, chromatin accessibility landscapes and GRNs, which we analyze together to highlight cell type-shared and -unique regulatory circuits. TFs were statistically ranked based on their ability to explain deuterosomal versus ciliated gene expression or signature genes shared by deuterosomal and ciliated cells (**Fig. 2A, Table S3C**). We also identify “TF modules”, composed of TFs with significantly overlapping regulatory interactions, that are predicted to co-regulate cellular processes (**Table S4**). Several of the top-ranked TF modules are predicted to operate in deuterosomal, ciliated or both cell types (**Fig. 2B**).

**Figure 2:**
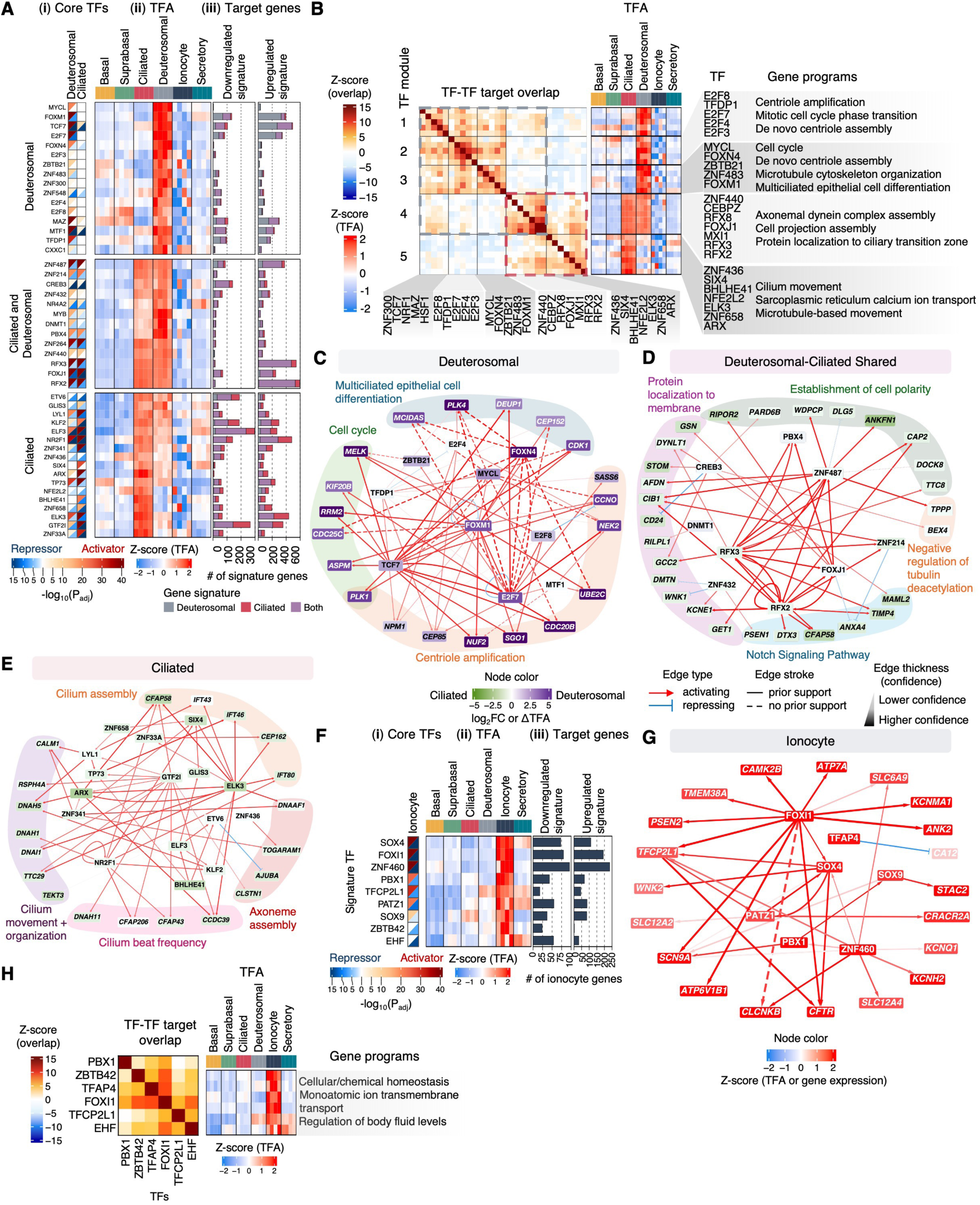
The GRNs governing deuterosomal cells, ciliated cells and ionocytes at baseline. (**A**) TFs predicted to regulate deuterosomal and/or ciliated gene programs at steady state, based on (**i**) a predicted activator role (enrichment of the TF’s activating gene targets in a cell type’s upregulated signature genes (red)) and/or a predicted repressor role (enrichment of repressed targets in a cell type’s downregulated signature genes (blue)), (**ii**) donor-resolved TFA estimates at steady state, (**iii**) number of shared and unique deuterosomal and ciliated cell type signature genes regulated per TF. (**B**) Five TF modules predicted to be active in ciliated and/or deuterosomal cells. The number of overlapping targets between TFs is indicated, and the TF modules are further annotated based on steady-state TFA estimates and the enrichment of their overlapping target genes in representative GO biological processes (Fisher’s exact test, P_adj_ < 0.05). Select regulatory interactions between core TFs and cell-type signature genes in (**C**) deuterosomal, (**D**) deuterosomal-ciliated shared and (**E**) ciliated cells. Nodes are colored by the difference in predicted TFA (TFs) or log_2_FC expression (target genes) between deuterosomal and ciliated cells at steady state. Edge color indicates activating (red) or repressing (blue) regulatory interactions, and thickness represents regulatory edge confidence. Some TFs are labelled in white for visual clarity. Genes are annotated based on GO biological process. (**F**) “Core” TFs governing Ionocyte gene signatures, with subpanels as in (**A**). (**G**) Select regulatory interactions between ionocyte “core” TFs and signature genes. Node color represents the z-scored TFA (TFs) or gene expression (targets) in ionocytes, relative to the other HAE cell types at steady state. (**H**) A TF module predicted to be active in ionocytes, annotated as in (**B**).

The top predicted regulators of deuterosomal-specific gene expression programs included TFs previously implicated in deuterosome processes (E2F7^52–54^, E2F4^65^, TFDP1^65^, FOXN4^6^) and others (FOXM1, MTF1, TCF7) (“deuterosomal” column of panel i, **Fig. 2A, 2C**). Many (e.g., FOXM1, TCF7, E2F7, MTF1) were predicted dual regulators, both activating and repressing deuterosomal-specific upregulated and downregulated gene signatures, respectively (panels i, iii, **Fig. 2A**). Consistent with deuterosome-mediated ciliogenesis pathways, E2F7 is predicted to cooperate with E2F4, E2F8 and TFDP1 (module 2, **Fig. 2B**) to regulate genes associated with centriole amplification (e.g., *CEP85*, *SASS6*, *CCNO*, *NEK2*, **Fig. 2C, Table S4B**). In addition to E2F family members, FOXM1 has established roles in cell-cycle progression^66–68^ and is a predicted deuterosomal regulator of cell cycle/centriole genes (e.g., *MELK*, *RRM2*, *ASPM*, *PLK1*, *CDC25C*, **Fig. 2C**).

Deuterosomal and ciliated cells shared several regulatory circuits, driven by MYB^64^, CREB3, ZNF432, PBX4 and others (**Fig. 2A, 2D**). Targets of these TFs are enriched in processes characteristic of these cell types^69^: cilium assembly (PBX4) and endoplasmic reticulum to Golgi vesicle-mediated transport (CREB3, ZNF432) (**Table S3**). A TF module, consisting of ZNF440, CEBPZ, FOXJ1, MXI1 and RFX family members, is predicted to coordinate dynein assembly and protein localization to the ciliary transition zone, processes in the intermediate stages of multiciliogenesis, initiated in deuterosomal cells and maintained in mature ciliated cells (module 4, **Fig. 2B**). While FOXJ1^58,70^ and MXI1^71^ have been previously established as regulatory factors in ciliated cells, both TFs also have high TFA in deuterosomal cells and regulate some deuterosomal (along with many ciliated cell) signature genes (**Fig. 2A**). Among ciliogenesis-associated RFX family members, RFX2 and RFX3 are predicted as the most likely regulators, due to higher expression (>100 transcripts per million (**TPM**)) and degree (i.e., number of target genes) than RFX5 and RFX8 (**Fig. S2C-D**).

We also identified TF regulators of gene expression programs unique to ciliated cells (“ciliated” column of panel i, **Fig. 2A, 2E**). While TP73 has established roles in ciliogenesis and ciliated cell maturation^72^, we identified additional TFs (e.g. ELK3, GTF2I, NR2F1, ZNF341) without previously described roles in ciliated cell processes (**Fig. 2A**). GTF2I is predicted to induce genes associated with cilium beat frequency and microtubule-based movement, while ELK3 is predicted to activate cilium assembly and structural genes, including *CFAP58*, *DNAAF1*, *IFT43*/*IFT46*/*IFT80*, *CEP162*, *RFX2* and *RFX3* (**Fig. 2E**, **Table S3**). Furthermore, several TFs, including ELK3, compose a TF module with high activity in ciliated cells, predicted to regulate genes involved in cilium and microtubule-based movement (module 5, **Fig. 2B**).

Taken together, our GRN analysis reveals a regulatory continuum between deuterosomal and ciliated cells. TFs predicted to regulate cell cycle and centriole biogenesis (“construction”) are more strongly associated with deuterosomal cells, whereas those regulating ciliary motility (“function”) are associated with ciliated cells. TFs controlling shared processes, such as assembly and transport machinery, are active in both. Thus, the GRN elucidated cell type-specific regulatory circuits in closely related cell types.

#### Ionocytes

Ionocytes are a rare, recently identified HAE cell type, with roles in epithelial ion transport and homeostasis^10,73,74^. In our GRN model, FOXI1, a TF established as necessary for ionocyte differentiation^10,47,74^, is a top-ranked ionocyte regulator (**Fig. 2F**), predicted to control essential ion transport genes integral to chloride transport and, consequently, mucosal hydration and airway clearance^73,74^ (e.g., *CFTR*, *SLC12A2*, *CLCNKB*, **Fig. 2G**). FOXI1, along with EHF, PBX1, TFAP4, TCFP2L1, and ZBTB42, compose a TF module with high TFA in ionocytes, predicted to coregulate chemical and cellular homeostasis, monoatomic ion transmembrane transport and regulation of body fluid levels (module 6, **Fig. 2H**).

#### Basal, suprabasal cells

As basal cells are the progenitor HAE cell types, many of the basal-specific regulators are stem cell-associated (e.g., TP63^75^, EGR3^76^, ZNF398^77^, **Fig. 3A, 3C**). In suprabasal cells, known differentiation TFs (e.g., KLF5^78^, FOSL1^79,80^, SREBF1^81^) are predicted to co-regulate epithelial cell differentiation programs, along with PRDM5, NKX2.1 and PLAGL1 (**Fig. 3B, 3D**). Cell-cycle regulators MYC, YBX1, SNAI2 and GLI2 coordinate the expression of basal-suprabasal shared gene signatures (**Fig. 3A, 3E**).

**Figure 3:**
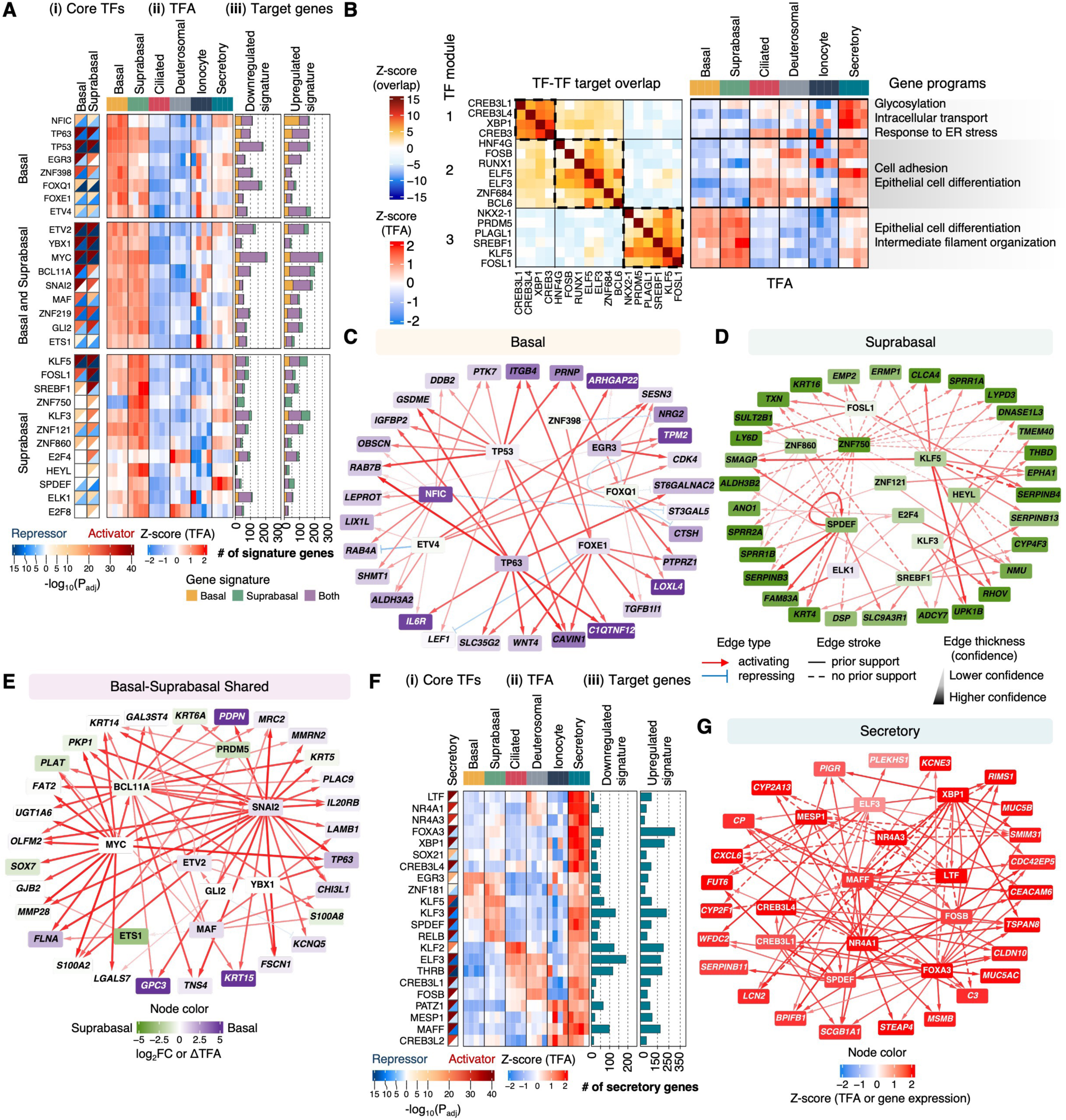
The GRNs governing basal, suprabasal and secretory cells at baseline. (**A**) “Core” TFs predicted to regulate basal and/or suprabasal gene programs at steady state, based on (**i**) predicted activator and/or repressor role in regulating cell type gene signatures, (**ii**) donor-resolved TFA estimates at steady state, (**iii**) the number of shared and unique basal and suprabasal signature genes regulated per TF. (**B**) Three TF modules predicted to be active in basal, suprabasal and/or secretory cells. The number of overlapping targets between TFs is indicated, and the TF modules are further annotated based on steady-state TFA estimates and the enrichment of their overlapping target genes in representative GO biological processes (Fisher’s exact test, P_adj_ < 0.05). Select regulatory interactions between core TFs and cell-type signature genes in (**C**) basal, (**D**) suprabasal and (**E**) shared basal-suprabasal cells. Nodes are colored by the difference in predicted TFA between basal and suprabasal cells (TFs) or log_2_FC expression (targets) at steady state. Edge color indicates activating (red) or repressing (blue) regulatory interactions. (**F**) “Core” TFs predicted to regulate secretory signature genes at steady state, annotated same as in (**A**). (**G**) Select regulatory interactions between secretory “core” TFs and signature genes. Node color represents the z-scored TFA (TFs) or gene expression (targets) in secretory cells, relative to the other HAE cell types at steady state.

#### Secretory cells

Core GRN analysis prioritized many known (e.g., XBP1^82,83^, FOXA3^84,85^, SPDEF^85^) and some novel secretory regulators (LTF, KLF3, MAFF, **Fig. 3F, 3G**). Secretory TFs compose two modules: one including XBP1 and CREB3 family members, predicted to regulate glycosylation, intracellular transport and ER stress response genes, and another, including ELF3, FOSB, and ELF5, which regulates cell adhesion and differentiation genes (**Fig. 3B, 3G**).

### IFNβ induces shared and cell type-associated gene expression responses in HAE

Having characterized the HAE cell types and their GRNs at steady state, we next explored their dynamic response to IFNβ. Focusing on scRNA-seq data from the five most abundant cell types, we identified 2,648 “interferon-responsive genes” (**IRGs**, FDR = 5%, |log_2_FC| > 1, **Table S5A**). IRGs exhibited both IFN-increased and IFN-decreased dynamics, clustering into shared and “cell type-associated” expression patterns (**Fig. 4A**, **Table S5B**). Shared “IFN-increased genes” (**IIGs**) constituted the largest cluster (548 genes, 21% of IRGs); these genes were similarly induced across all HAE cell types analyzed, with maximal expression at either 2 or 6h. Shared IIGs were enriched for canonical interferon-stimulated genes (**ISGs**^37^) and ISGF3 target genes as predicted by our GRN (Fisher’s exact test, P_adj_ < 10^-100^, P_adj_ < 10^-116^, respectively). A small cluster of “IFN-decreased genes” (**IDGs**, 232 genes, 9% of IRGs) was equivalently downregulated across cell types in response to IFNβ.

**Figure 4:**
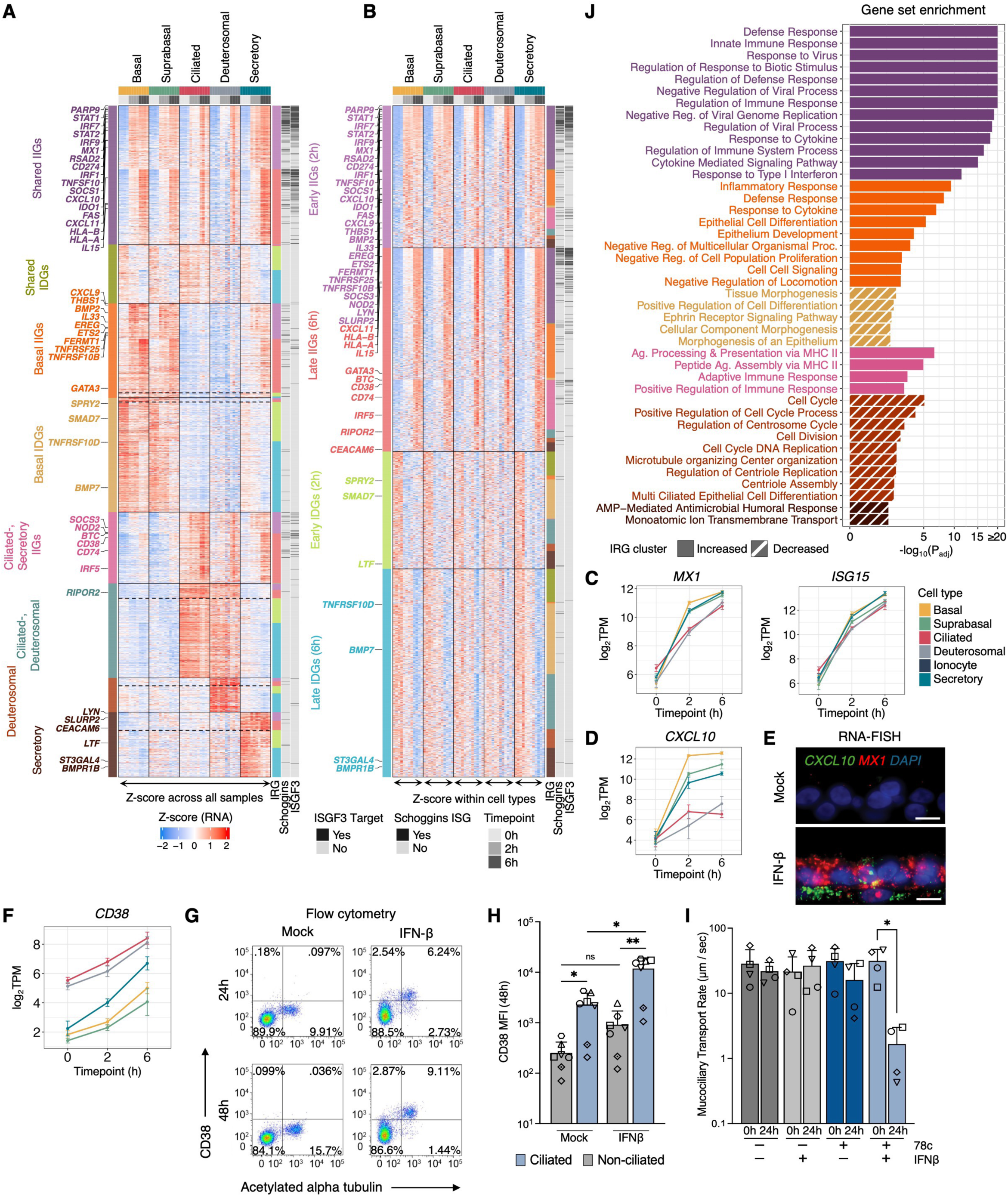
Shared and cell-type-associated transcriptional responses to interferon in the HAE. (**A-B**) Pseudobulk gene expression of 2,648 interferon-regulated genes (**IRGs,** FDR = 5%, |log_2_(FC)| > 1; n = 4 donors). Genes were (**A**) z-scored across all pseudobulk samples or (**B**) z-scored *within* each cell type (to normalize away cell-type differences in gene expression and enhance visualization of dynamic trends within each cell type). Dotted lines in (**A**) distinguish IIG and IDG subclusters based on (**B**). Mean expression (log_2_(transcripts per million)) of (**C**) shared (*MX1*, *ISG15*) or (**D**) cell type-associated (*CXCL10*) IRGs across cell types and time points; error bars represent standard error. (**E**) RNA fluorescent *in situ* hybridization of HAE culture cross-sections. (**F**) *CD38* expression, presented as in (**C-D**). (**G**) Flow cytometry measurement of CD38 and acetylated alpha tubulin (ciliated cell marker) expression in HAE cells treated with IFNβ for 24 or 48h. (**H**) Mean fluorescence intensity of CD38 in acetylated alpha tubulin-negative (i.e. non-ciliated) or -positive (ciliated) cells. Mean with standard deviation across five experimental replicates that utilized 4 HAE donors is shown (unpaired t-test with Welch’s correction, *P < 0.05, **P < 0.01, ns = not significant). (**I**) Mucociliary transport rate in mock or IFNβ-treated HAE cultures in the presence or absence of CD38 inhibitor (78c). Data include four experimental replicates that utilized 3 HAE donors. The mean (with standard deviation) of the median MCT values for each culture are shown (paired t-test, *P < 0.05). Cultures from the same donor within an experimental replicate are indicated by symbols in both (**H**) and (**I**). (**J**) Select enriched GO biological processes for clusters in (**A**). For clusters with mixed IIG/IDG dynamics (e.g., ciliated-deuterosomal), IIGs and IDGs were considered separately and are annotated with bar shading (solid for IIGs, dashed for IDGs). Fisher’s exact test, P_adj_ < 0.05.

In contrast to a general paradigm in which all cell types exhibit similar gene expression dynamics following IFN stimulation, surprisingly, most IRGs (70%, 1,868 genes) varied dramatically in their induction amplitude across the HAE cell types. In general, these “cell type-associated” IRGs were only differentially expressed in particular HAE subsets and differed in their initial, steady-state abundances (**Fig. 4A**). Several of the clusters were driven by cell type and exhibited coordinated up- or downregulation to IFN: “basal IIGs”, “basal IDGs” and the “ciliated-secretory IIGs”. In contrast, the three remaining clusters were dominated by cell type-associated expression and exhibited a mix of IFN-increased and -decreased genes: “ciliated-deuterosomal”, “deuterosomal” and “secretory”.

To better visualize IRG time-course dynamics, we standardized gene expression within cell types, thereby normalizing cell type-associated differences in gene expression values (**Fig. 4B**). With this data scaling strategy, IRGs segregated into four groups: “early” IIGs (maximal at 2h), “late” IIGs (maximal at 6h) and analogously categorized early and late IDGs. This multi-level view of the gene expression responses (**Figs. 4A-B**) revealed that, while the steady-state and IFN-responsive expression levels of the cell type-associated IRGs differed across HAE cell types, time dynamics were largely consistent. Early IIGs had the strongest enrichment in canonical ISGs (OR=16.3, P_adj_ <10^-16^) and predicted ISGF3 targets (OR=128.2, P_adj_ <10^-16^), but late IIGs were also enriched for canonical ISGs (OR=6.0, P_adj_ <10^-16^) and for ISGF3 targets (OR=16.2, P_adj_ <10^-16^). ISGF3, STAT1 and IRF family members (IRF1, IRF5, IRF7) are predicted as regulators for 50% of IIGs but for only 8% of IDGs. Thus, the complex transcriptional response following IFN stimulation of HAE cannot be fully attributed to canonical IFN-responsive TFs directly downstream of IFNAR/JAK/STAT signaling.

Cell type-associated IRG expression dynamics were apparent for all HAE cell types. For example, among well-known ISGs, *MX1* and *ISG15* exhibited consistent induction across cell types (**Fig. 4C**), while *CXCL10* was more highly induced in basal, suprabasal and secretory cells relative to ciliated and deuterosomal cells (**Fig. 4D**). We confirmed the cell type-associated expression of *CXCL10* transcripts by RNA fluorescence *in situ* hybridization (**RNA-FISH**). While *MX1* transcripts were induced throughout the pseudostratified epithelium in response to IFNβ, *CXCL10* signal was localized primarily to the basolateral aspect of the culture, consistent with basal cell-associated induction (**Fig. 4E**). CXCL10 promotes chemotaxis of immune cells in inflammatory and antiviral contexts^86^. IFNβ-dependent *CXCL10* expression by basal cells would place the strongest signal proximal to blood vessels, consistent with a role in leukocyte recruitment.

As another example, although *CD38* is upregulated in response to IFN across HAE cell types, *CD38* is upregulated at steady-state and following IFN stimulation in ciliated and deuterosomal cells, relative to other cell types (**Fig. 4F**). Flow cytometry confirmed higher steady-state and IFN-induced CD38 protein expression in acetylated α-tubulin positive (i.e. ciliated) cells (**Fig. 4G-H**). In immune cells, CD38 has been implicated in immune regulation, calcium homeostasis, oxidative stress response and airway remodeling and repair^87^, but the functional consequence of its IFNβ-induced expression in ciliated cells is unknown. Given the important role of ciliated cells in propelling mucus, we assessed the potential impact of CD38 on mucociliary transport (**MCT**). Pharmacological inhibition of CD38 significantly reduced MCT in IFN-treated, but not steady-state HAE (**Fig. 4I**), providing initial evidence that cell type-associated IRGs may contribute to HAE tissue-level processes in the context of the innate antiviral response.

To predict the potential functional consequences of cell type-associated IFN-responsive gene expression, we performed gene set enrichment testing for each cluster in **Fig. 4A** (**Fig. 4J**, **Table S5C**). We identified 317 GO terms enriched in the shared IIG cluster, and 168 GO terms enriched across all cell type-associated IRG clusters. As expected, GO terms associated with shared IIGs involved well-established and overlapping innate immune response gene sets, including canonical ISGs such as *ISG15*, *ISG20*, *OAS1*, *IFIT1* and *IFI44*. Basal IIGs were associated with inflammatory and defense responses, negative regulation of locomotion, and negative regulation of cell population proliferation. The upregulation of genes inhibiting proliferation and differentiation was matched by IFN-induced downregulation of “positive regulation of cell differentiation” and “morphogenesis of an epithelium” processes in basal cell (basal IDGs). Ciliated/secretory IIGs were enriched for MHC Class II antigen presentation genes, while secretory cell downregulated genes were involved in monoatomic ion transmembrane transport and antimicrobial peptide-mediated antimicrobial responses.

Thus, in addition to “classic” antiviral effector gene programs, IFN signaling in HAE induces cell type-associated gene expression responses that modulate different biological processes, consistent with cell type function and the potential for specialized roles in the antiviral response.

### IFNβ induces shared and cell type-associated chromatin accessibility responses in HAE

We next investigated how the chromatin landscape might contribute to shared and cell type-associated IRG expression patterns. Given the higher dimensionality and greater sparsity of snATAC-seq^88^, we limited our analysis of IFN-dependent accessible chromatin to the four most abundant HAE cell types, identifying 6,210 interferon-responsive accessible chromatin regions (**Fig. 5A-B**, **S3A-B**, **Table S6**). Mirroring IRGs, most of the IFN-induced accessibility chromatin regions exhibited cell type-associated dynamics. However, chromatin accessibility dynamics differed from gene expression dynamics in several ways. While IFN transcriptionally induced comparable numbers of up- and down-regulated genes, IFN mainly induced chromatin opening (97% of differential peaks). While gene expression exhibited both early and late dynamics (peaking in amplitude at 2 or 6h, **Fig. 4B**), IFN-responsive chromatin accessibility peaked at 2h (**Fig. 5A-B**). These observations are consistent with chromatin accessibility preceding gene expression changes. Indeed, the RNA expression of genes proximal to IFN-increased accessible chromatin regions peaked at 6h (**Fig. 5C**).

**Figure 5:**
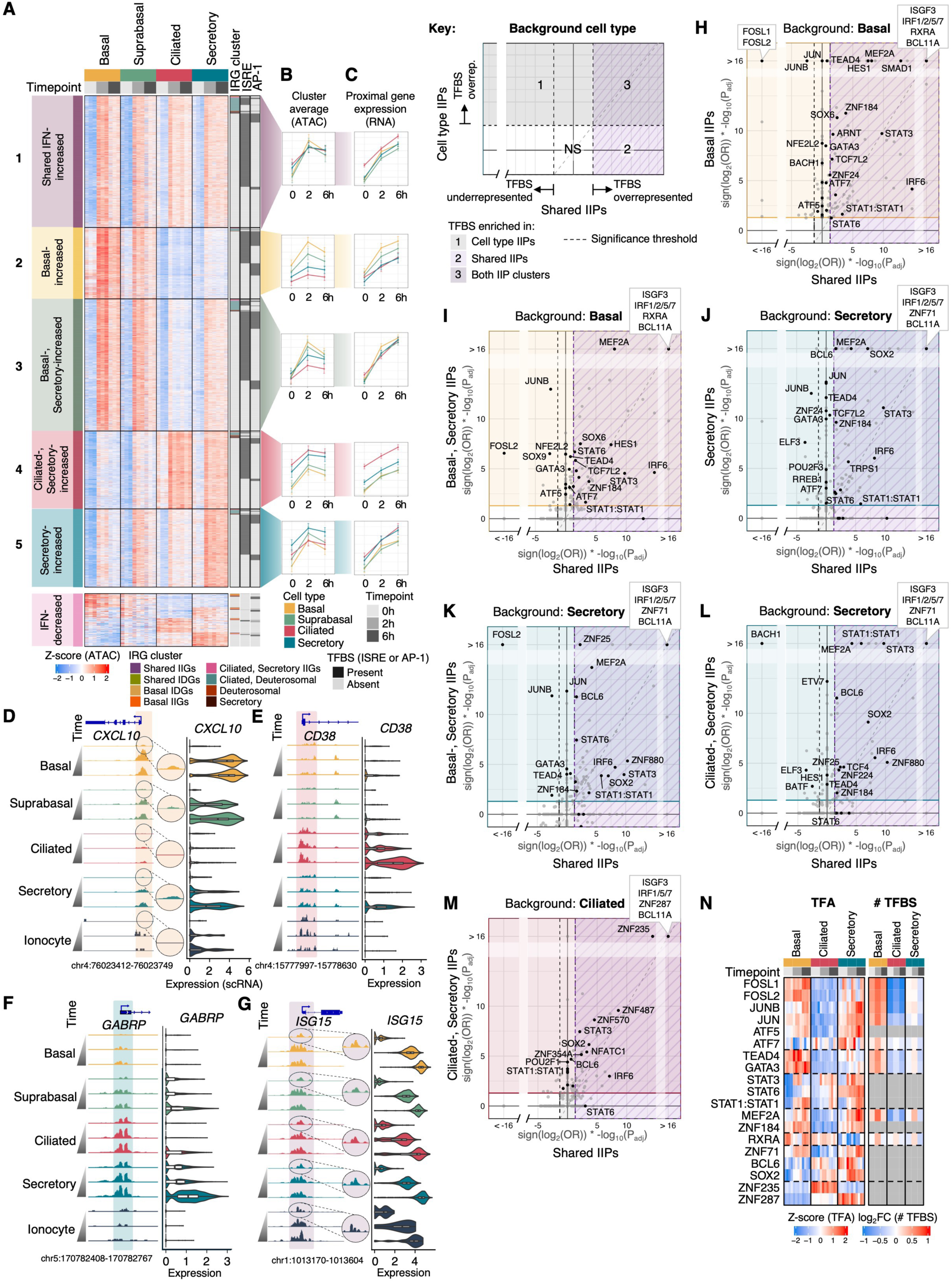
Shared and cell type-associated TFs are associated with interferon-dependent chromatin accessibility. (**A**) Pseudobulk chromatin accessibility of interferon-responsive peaks (P_adj_ < 0.05, |log_2_(FC)| > 1, n = 3 donors). Each peak is annotated (right-hand side) based on proximity (± 2kb TSS) to an IRG and colored according to the IRG clusters in Fig. 4A, occurrence of an ISRE motif or *maxATAC*-predicted AP-1 TFBS (considering FOS, FOSL1, FOSL2, JUN, JUNB, JUND, ATF3, ATF7 model predictions in any cell type-time point simulated). (**B**) Mean peak accessibility per cluster; error bars are standard error of cluster means across biological replicates. (**C**) Mean gene expression (log_2_(TPM)) of any gene proximal (±2kb TSS) to a peak within each cluster; error bars as in (**B**). Chromatin accessibility dynamics at IRGs, with zoom-in visualizations of steady-state accessibility at (**D**) *CXCL10*, (**E**) *CD38*, (**F**) *GABRP* and (**G**) *ISG15*; ATAC signal was normalized per million reads. Violin plots of distribution of gene expression (log-normalized UMIs per 10k UMIs). (**H**-**M**) The enrichment of TFBS in cell type-associated (y-axis) or shared (x-axis) interferon-increased peak clusters (from (**A**)), where over- or under-enrichment are indicated by positive or negative sign, respectively, based on multiplication of log_2_(odds ratio) and -log_10_(P_adj_). To control for baseline accessibility in the corresponding cell type, p-values were estimated via simulation (**Methods**). TFs with concordant TFA and TFBS enrichments are indicated with black dots and a select subset are labeled. IIP = IFN-increased peak, NS = not significant. (**N**) For select TFs, TFA estimates and the predicted number of TFBS (limited to TFs with *maxATAC* models available). The number of TFBS is presented as log_2_(FC) relative to the mean number of TFBS across timepoints and cell types.

In total, we identified five IFN-increased peak clusters: one with HAE-shared dynamics, two with maximal accessibility in basal and suprabasal cells, and two with maximal accessibility in ciliated or secretory cells (**Fig. 5A**). The shared IFN-increased peaks (cluster 1), identified from analysis of the most abundant cell types, also exhibited chromatin opening trends in rare HAE cell types (ionocytes and NE cells, **Fig. S3C-D**). Peaks with cell type-associated dynamics were enriched proximally to genes with corresponding expression patterns (**Fig. S3E**). However, relative to all accessible chromatin regions detected in our study, the IFN-dynamic peaks were more distal to gene promoters (P < 10^-16^, K-S test, median distance of 21kb vs. 13kb to the nearest TSS, **Fig. S3F**). Consistent with the role of enhancers in regulation of cell-type specificity, the cell type-associated dynamic peaks were more promoter-distal than peaks with shared dynamics (P < 10^-16^, K-S test, median distance to TSS of 25kb vs. 12kb, **Fig. S3G**).

Peak clusters with cell type-associated opening dynamics exhibited priming (i.e., increased accessibility at steady state in the corresponding cell type, **Fig. 5B**, **S3H**). As an individual example, chromatin proximal to the *CXCL10* promoter had elevated accessibility in basal cells at steady state, which corresponded to greater IFN-dependent chromatin opening and greater *CXCL10* induction in basal cells (**Fig. 5D**). Analogously, ciliated/secretory cell-associated IRG *CD38* and secretory cell-associated IRG *GABRP* exhibited cell type-associated chromatin priming (**Fig. 5E-F**). In contrast, for the shared IRG *ISG15*, all cell types exhibited similar steady-state accessibility at the *ISG15* promoter, which corresponded with its uniform mRNA induction across cell types upon IFN stimulation (**Fig. 5G**).

Whether cell type-associated or HAE-shared, each cluster of IFN-increased accessible chromatin regions was highly enriched for IFN-stimulated response elements (**ISREs**, Fisher’s exact test, P < 10^-16^), suggesting that TFs beyond STAT1, STAT2 and IRF family members contribute to the cell type-associated chromatin dynamics in response to IFN (**Fig. 5A**). To distinguish TFs driving cell-type-associated chromatin accessibility dynamics from TFs that were merely cell type-associated at baseline, we designed a TFBS enrichment analysis that controlled for baseline accessibility in each cell type (**Methods**). For example, to identify TF drivers of the basal-associated IFN-increased ATAC-seq peak cluster, the number of TFBS occurrences in that cluster was compared to background peaks with matched baseline accessibility in basal cells. Furthermore, we required that candidate TFs exhibit IFN-dependent protein activity in the corresponding cell type (TFA, as estimated by the GRN, or number of TFBS, as predicted by *maxATAC*^44^). Thus, our final set of candidate IFN-dependent chromatin-accessibility regulators was supported by both enrichment of TFBS and TFA estimates (**Fig. 5H-M**, **S4A-C, Table S6**).

As anticipated from our initial analysis (**Fig. 5A**), ISGF3 and IRF2/5/7 were top candidate regulators of IFN-dependent chromatin opening across clusters. The analysis also nominated cell type-specific TF drivers of chromatin accessibility, some with binding sites specifically enriched in cell-type associated chromatin regions while others with binding sites enriched in both shared and cell-type associated peaks (**Fig. 5H-M**). For example, in basal cells, several AP-1 factors exhibited interferon-increased activities (FOSL1, FOSL2, JUNB, **Fig. 5N**), and their binding sites were uniquely enriched (FOSL1, FOSL2, JUN, JUNB, ATF5, ATF7) in basal cell-associated IFN-induced chromatin opening (clusters 2, 3) but not HAE-shared IFN-induced opening regions (cluster 1) (**Fig. 5H-I**). Similarly in secretory cells, JUNB and JUN activities were IFN-increased (**Fig. 5N**), and the binding sites of several AP-1 factors (JUN, JUNB, FOSL2, ATF7) were enriched in basal-secretory (cluster 3) or secretory-associated (cluster 5) (**Fig. 5J-K**) but neither the HAE-shared (cluster 1) nor ciliated-secretory (cluster 4) IFN-increased peaks (**Fig. 5L**). In ciliated cells, AP-1 factors were generally lowly expressed, predicted to have fewer binding sites genome-wide (**Fig. 5N**, panel v**, S4A-C**) and their binding sites did not colocalize in the ciliated-associated, IFN-induced peaks (cluster 4) (**Fig. 5M**). Collectively, AP-1 factors were implicated in IFN-induced chromatin opening specific to basal and secretory but not ciliated cells. Several other TFs exhibited similar basal- and secretory-associated binding patterns, including TEAD4 and GATA3 (**Fig. 5N**).

Beyond ISGF3, many TFs were associated with both cell-type-associated and shared IFN-induced chromatin accessibility. For example, across all cell types, STAT1:STAT1, STAT3 and STAT6 had binding sites enriched across the IFN-increased peaks, with corresponding TFA dynamics (**Fig. 5H-N**). In secretory and basal cells, MEF2A, STAT6 and ZNF184 were associated with both shared and secretory- and basal-associated peak clusters (**Fig. 5H-L, N**), while RXRA was a top-ranked basal-specific regulator of basal-associated and shared IFN-increased peaks (**Fig. 5H-I**) and ZNF71 was a top-ranked secretory-specific regulator of secretory-associated and shared peaks (**Fig. 5J-L, 5N**). In ciliated and secretory cells, BCL6 and SOX2 were predicted regulators of IFN-induced chromatin accessibility (**Fig. 5J-N**). Finally, unique to ciliated cells, the binding sites of ZNF235 and ZNF287 were highly enriched in both shared and cell-type associated chromatin opening (**Fig. 5M-N**).

Taken together, our analyses highlight known regulators of IFN-induced chromatin accessibility but also identify additional TFs predicted to drive the shared and cell type-associated chromatin dynamics observed in the HAE response to IFN.

### Dynamic and cell type-associated TFs finetune the IFN responses of HAE cell types

We next used the GRN to identify TF regulators orchestrating the IFN-induced gene expression changes. We hypothesized that the cell type-associated IFN-induced gene expression patterns were due to chromatin priming and differences in IFN-dependent TF activities across the HAE cell types. Several lines of evidence supported chromatin priming: The cell type-associated IRG clusters (**Fig. 4A**) exhibited increased baseline accessibility in the corresponding cell types, and, for IDGs only, increased gene expression in the corresponding cell type (**Fig. 6A**). Thus, at least some of the TFs that maintain HAE cell type identities at steady-state (**Fig. 1J**) are implicated in IFN-induced gene expression responses via priming, even if IFN stimulation does not alter the protein activities of these “baseline” TFs. However, mirroring our analysis of IFN-induced chromatin dynamics (**Fig. 5**), we also sought to identify TFs whose IFN-responsive activities drove gene expression dynamics.

**Figure 6:**
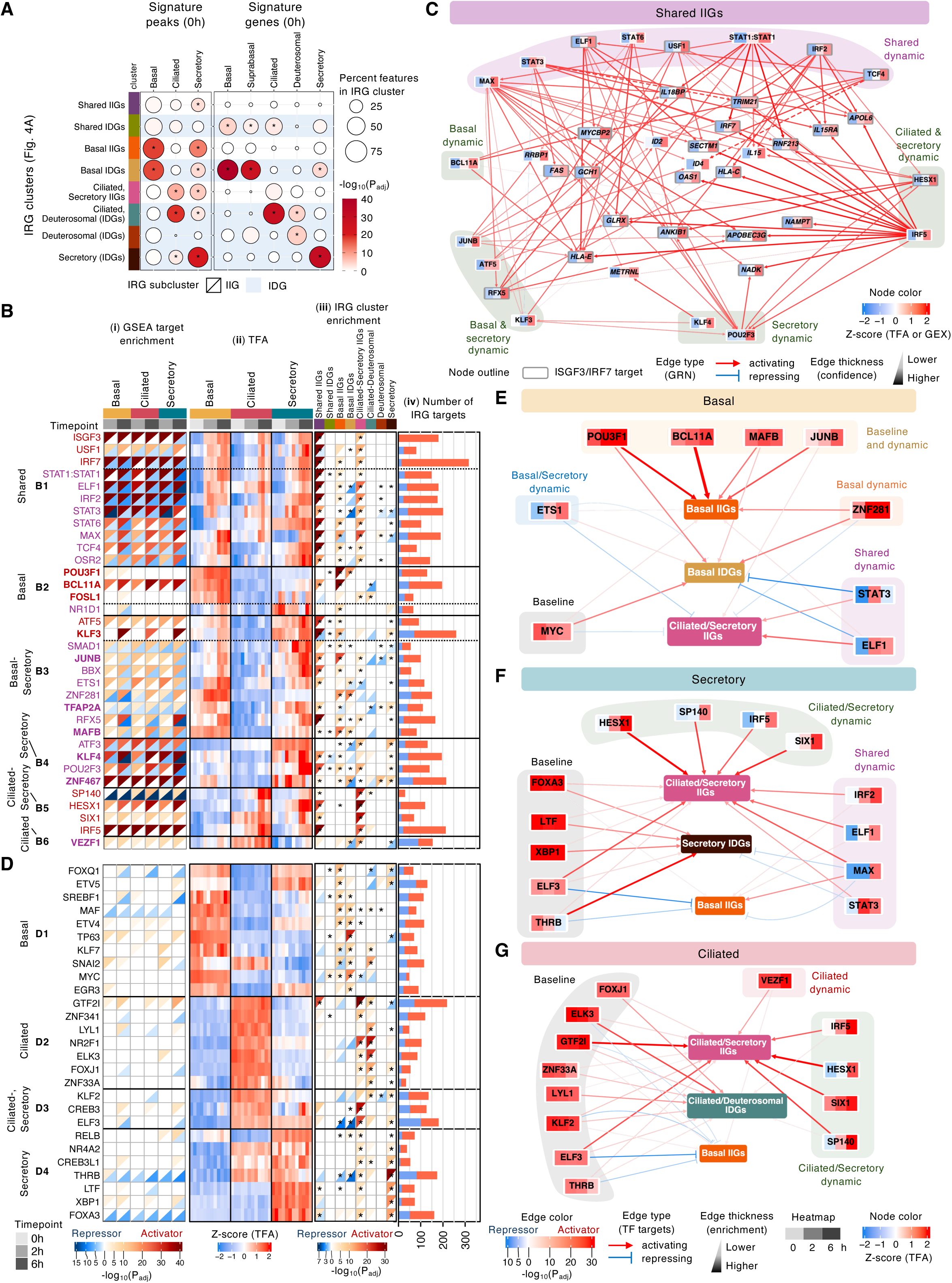
Cell type-associated IRGs are regulated by baseline and interferon-responsive TFs. (**A**) Enrichment of IRG clusters from Fig. 4A in genes proximal (±2kb of TSS) to cell-type signature peaks (left) or signature genes (right), derived from steady state (i.e., from Fig. 1C**-D**). For IRG clusters containing both IIGs and IDGs, the enrichment was limited to the dominant IIG or IDG pattern, and predominantly IDG clusters are highlighted with a light blue strip (Fisher’s exact test, *P_adj_ < 0.05). (**B**) Data supporting the top-predicted **“**dynamic” TF regulators of IRG patterns, a subset (bolded text) are additionally “baseline” TFs, predicted to prime chromatin proximal to cell type-associated IRGs at steady state. We report (**i**) “within-cell type” predicted activator role (red: enrichment of the TF’s activating gene targets in a cell type’s IFN-upregulated genes at 2 or 6h relative to 0h) and/or a predicted repressor role (blue: enrichment of the TF’s repressed gene targets in a cell type’s IFN-downregulated genes at 2 or 6h relative to 0h), (**ii**) TFA dynamics across cell types and time points, (**iii**) potential role in priming of the IRG clusters: asterisks indicate that TFBS (from *maxATAC* or motif scanning) are enriched proximally to genes within the IRG cluster; Fisher exact test, P_adj_ < 0.05; while box shading indicates predicted activator or repressor role, based on TF’s target enrichments in the IIGs or IDGs of each IRG cluster, and (**iv**) the number of activating or repressing IRG targets per TF. (**C**) Select regulatory interactions between shared or cell type-associated dynamic TFs and shared IIGs. Node colors represent the z-scored gene expression (target nodes) or TFA (TF nodes) over time, averaged across basal, ciliated and secretory cells. TFs are grouped based on their cell type-associated activities and dynamics. (**D**) Top baseline TF regulators of cell type-associated expression of IRG clusters (see **Methods** for inclusion criteria), annotated as in (**B**). (**E**-**G**) Network diagrams showing the regulation of IRG clusters by baseline and/or IFN-responsive dynamic TFs. TFs are shaded according to z-scored TFA (donors averaged per time point) in the represented cell type. Edges are colored based on the adjusted p-value of TF target enrichment from panel iii in **6B, D**.

We identified IFN-responsive TFs per cell type based on (1) IFN-increased TFA and (2) enrichment of TF target genes among IRGs. To capture IFN-responsive TFs (as opposed to nondynamic steady-state TFs involved in priming), IRGs for enrichment analysis were defined per cell type relative to that cell type’s baseline (e.g. 2h v 0h in basal cells), so that steady-state effects (due to cell type) were normalized. Although the cell type-associated IRGs were more numerous than the shared IFN-increased or decreased genes (**Fig. 4A**), the enrichments of STAT1, STAT2 and IRF TF targets among IRGs in basal, secretory or ciliated cells at either time point (2h or 6h relative to baseline) were so strong that we performed a second round of analysis, omitting genes from both the shared IIG and shared IDG clusters (**Fig. 4A**), to increase signal for TFs driving cell type-associated IRGs. ISGF3, IRF7, STAT1:STAT1 targets were still highly enriched in the IRGs per cell type, even upon removal of the shared cluster genes, but we additionally prioritized 34 IFN-responsive TFs with targets enriched in basal-, secretory- or ciliated-cell IRGs (**Fig. 6B**). These TFs were categorized as shared or cell type-associated, based on IRG enrichment (panel i) and TFA (panel ii), and annotated as activators, repressors or both, based on (1) total number (panel iv) and enrichment of activated or repressed target genes in IIGs or IDGs (panel i). We also implicated TFs in the regulation of IRG clusters from **Fig. 4A**, based on enrichment of proximal TFBS and activating or inhibiting interactions (panel iii).

As an example, we identified 11 TFs with cell type-shared IFN-responsive activities (cluster B1, **Fig. 6B**), which, along with some cell type-associated dynamic TFs (e.g., BCL11A, KLF3, etc., from clusters B2-5), have gene targets highly enriched in shared IIGs (panel iii, column “Shared IIGs”, **Fig. 6B**). These shared IIG regulators and select target genes are highlighted in **Fig. 6C**, where, for legibility, the numerous ISGF3 target genes are outlined in charcoal. Cell type-shared versus cell type-restricted TFA was not the only source of variability among the shared-IIG regulators. For example, ISGF3 and IRF7 were predicted to be mainly activators, as supported by both their IRG cluster enrichment analyses (panel i, iii) and number of activating IRG targets (panel iv, **Fig. 6B**). In contrast, STAT3 and recently IFN response-implicated regulator ELF1^89^ were both predicted to activate IIGs and repress IDGs (panels i, iii), suggesting dual activities. The shared IIG regulators also varied in their predicted dynamics: Many noncanonical TFs (e.g., KLF3, STAT6, MAX, TCF4, POU2F3, and HESX1) had greater enrichment of targets in the 6h IRGs (6h v. 0h comparison) as opposed to 2h IRGs, suggesting a delay in activity, while, as expected, direct mediators of IFN signaling (e.g., ISGF3) were highly enriched in both 2h and 6h IRGs (panel i, **Fig. 6B**).

Baseline TFs were defined as core TFs (elevated TFA and drivers of steady-state cell type identities, e.g., **Fig. 1J**) that were likely to contribute to accessible chromatin priming of cell-type-associated IRGs. We ensured that the baseline TFs had (1) TFBS enriched proximally to and (2) positive or negative GRN targets enriched in IIGs or IDGs of the corresponding cell type (as shown in panel iii, **Fig. 6C, S4D-F**). We report 27 baseline TFs, of which ten (bolded in **Fig. 6B**) were also IFN responsive. The remaining baseline TFs lacked IFN-increased TFAs (panel ii) and explained IRG differences across cell types (panel iii) but not within cell types (i.e., weak or no enrichments in panel i, **Fig. 6D**).

While the dynamic TFs were sufficient to explain the shared IIG expression patterns (**Fig. 6C**), the explanation of cell type-associated IFN response genes required both baseline and dynamic TFs (**Fig. 6E-G**). In each cell type (basal, secretory, ciliated), we nominated regulators driving expression of the cell type’s associated IIGs and IDGs as well as regulators disabling inappropriate expression of “alternative-cell type” IIGs. For example, in basal cells, baseline TFs (POU3F1, BC11A, MAFB, JUNB, MYC) had TFBS enriched proximally to basal IIGs and basal IDG clusters, setting elevated baseline accessibility and gene expression at 0h (**Fig. 6E, S4D**). The TFAs of several baseline TFs increased upon IFN stimulation (POU3F1, BC11A, MAFB, JUNB), and, in concert with other basal-associated dynamic TFs (ETS1, ZNF281), are predicted to activate basal IIGs. In contrast, the IFN-increased TFAs of the shared dynamic TFs STAT3 and ELF1 are predicted to repress basal IDGs. The decreased TFA of baseline TF MYC, an activator of basal IDGs, is also predicted to contribute to the downregulation of basal IDGs. Finally, MYC and ETS1 are predicted to repress inappropriate expression of ciliated/secretory IIGs, and ETS1 has TFBS enriched proximal to these genes in basal cells (**Fig. S4D**).

A similar model emerges in secretory cells (**Fig. 6F**). Baseline TFs (FOXA3, LTF, XBP1, ELF3, THRB) have TFBS enriched proximally to ciliated/secretory IIGs and secretory IDGs and are predicted to set elevated chromatin accessibility at and expression of these genes in secretory cells at steady state. Induction of ciliated/secretory IIGs requires activation from the ciliated/secretory dynamic TFs (HESX1, SP140, IRF5, SIX1) and shared dynamic TFs (IRF2, ELF1, MAX, STAT3). MAX and STAT3 have predicted dual activities and are predicted drivers of secretory IDG dynamics, via repressive interactions. Diminution in THRB TFA, an activator of secretory IDGs, is also predicted to contribute to secretory IDG dynamics. Finally, MAX, ELF3 and THRB are predicted to repress “alternative-cell type” basal IIGs, with support from ELF3 TFBS, enriched proximally to the basal IIGs in secretory cells (**Fig. S4E**).

In ciliated cells, a combination of baseline and dynamic TFs are also predicted to orchestrate ciliated/secretory IIGs, ciliated/deuterosomal IDGs and repress inappropriate basal IIG expression (**Fig. 6G**). The uniquely ciliated-associated dynamic baseline TF VEZF1 collaborates with the ciliated-secretory dynamic TFs (e.g., HESX1, SIX1) to induce ciliated/secretory IIGs. While our analysis nominated repressors (with IFN-increased TFAs) and activators (with IFN-decreased TFAs) of cell type-associated IDGs in basal and secretory cells, reduction of TFA for activating TFs (ZNF33A, THRB) was the only high-confidence predicted mechanism driving ciliated/deuterosomal IDG dynamics in ciliated cells. Like secretory cells, ELF3 and THRB were predicted repressors of inappropriate basal IIG expression, along with ciliated-specific KLF2.

Taken together, GRN analysis supports a regulatory logic in which baseline, lineage-determining TFs prime the chromatin landscape and establish steady-state expression of cell type-associated IRGs, and a partially overlapping set of IFN-responsive TFs coordinate the observed transcriptional dynamics alongside canonical ISGF3/IRFs. This interplay establishes both a common antiviral state and the cell type-associated IFN responses.

### Prediction of gene regulatory mechanisms underlying cell type-associated IFN responses

Based on these data, we took a global approach, connecting regulators in our working model (**Fig. 6B-G**) to potential cell type-associated IRG phenotypes (GO biological processes in **Fig. 4J**). In total, 61 baseline and/or dynamic TFs had gene targets enriched in GO terms (**Fig. S5**). Next, we explored the gene regulation of mucin and chemokine genes, as both gene sets exhibited a diversity of dynamic and cell type-associated gene expression patterns (**Fig. 7**).

**Figure 7:**
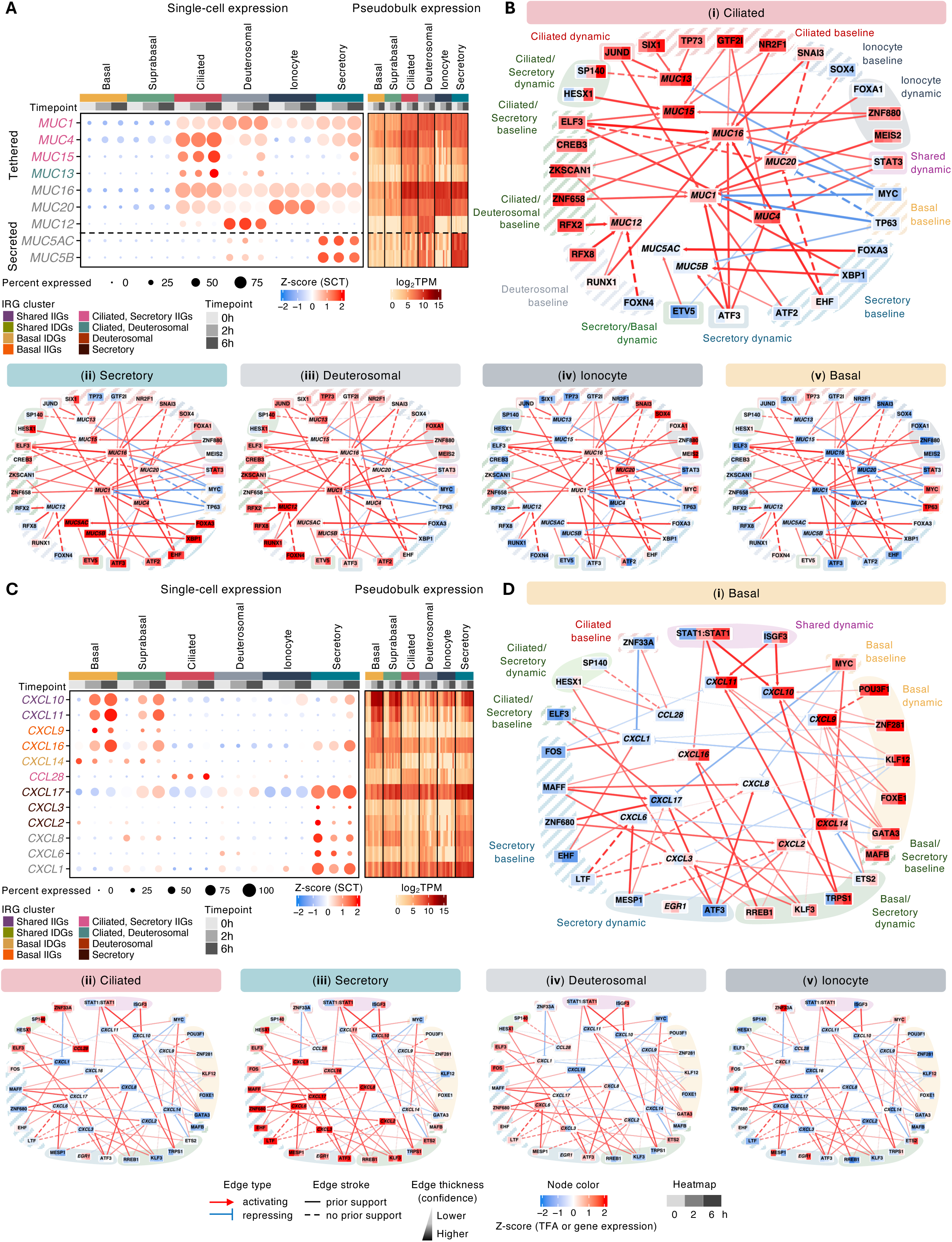
Cell type-associated mucin and chemokine gene regulatory networks. (**A**) Gene expression of mucins, normalized using *SCTransform* (dot plot) or log_2_(TPM) (heatmap). Mucins are colored according to IRG cluster membership or gray, if non-dynamic. (**B**) Select regulatory interactions between shared or cell type-associated TF regulators and mucin genes. Nodes colors represent the z-scored TFA (TFs) or gene expression (targets) in each cell type. Edge color and thickness represent the GRN-predicted edge confidence. TFs are grouped based on their cell type-associated activities and dynamics. (**C**) Gene expression of chemokine genes, annotated as in (**A**). (**D**) Select regulatory interactions describing chemokine regulation, annotated as in (**B**).

We detected nine mucins in our dataset (**Fig. 7A**). While the secreted mucins (*MUC5AC, MUC5B*) were constitutively expressed by secretory cells, tethered mucin expression was often dynamic and varied across ciliated, deuterosomal, ionocyte and secretory cells. Among IFN-responsive tethered mucins, *MUC1/4/15* clustered with cell type-associated “Ciliated, Secretory IIGs”, while *MUC13* was a member of the “Ciliated, Deuterosomal” IRG cluster. On average, ciliated cells had the highest expression of tethered mucins. In each cell type, we visualized the dynamics of predicted regulators and their mucin targets (**Fig. 7B**). At steady-state, ciliated expression of tethered mucins was supported by ciliated-specific TFs (JUND, SIX1, GTF2I, NRF2) and ciliated-secretory-shared TFs (ZKSCAN1, ZNF658, ELF3). Upon IFN stimulation, the shared dynamic regulator STAT3 was a predicted activator of *MUC1*/*4* expression, while ciliated/secretory or ciliated dynamic TFs SP140, HESX1 and JUND contributed to *MUC13/15* induction in ciliated cells (panel i, **Fig. 7B**). In secretory and deuterosomal cells, ATF3 exhibits interferon-increased TFA in both cell types and is predicted to drive *MUC1/4* expression (panels ii-iii). Secretory TFs FOXA3, XBP1, ATF3 and ETV5 are predicted to maintain high stead-state expression of the secreted mucins (*MUC5AC*, *MUC5B*) and/or tethered mucins (*MUC1*, *MUC16*). Finally, SOX4 and SNAI3 were predicted to activate the ionocyte-associated tethered mucin *MUC20* (panel iv), while RUNX1 and ZKSCAN1 explain high constitutive expression of *MUC12* by deuterosomal cells (panel iii). With the exception of MUC1, the potential immunomodulatory roles of the cell type-associated, IFN-induced mucins during viral infection are unknown^90,91^.

Many inflammatory stimuli, including IFN-β, induce the expression of chemokines^92^. Here, we detected twelve chemokines, of which nine were dynamic (**Fig. 7C**). Induction of chemokines *CXCL9/10/11/16* was strongest in basal cells, with weaker response dynamics in suprabasal and/or secretory cells. *CXCL17* clustered with secretory IRGs, due to high steady-state expression in secretory cells, but was also IFN-increased in basal and suprabasal cells. Only one chemokine, *CCL28*, was a ciliated-associated IIG. The three remaining chemokines were IDGs in basal (*CXCL14*) or secretory cells (*CXCL2/3*). We queried the GRN for underlying regulatory mechanisms (**Fig. 7D**). In basal cells, basal baseline and dynamic TFs (POU3F1, GATA3, FOXE1) collaborated with shared (ISGF3, STAT1:STAT1) and basal/secretory (TRPS1) dynamic TFs to upregulate *CXCL9/10/11* more strongly than secretory cells (panel i). In contrast, *CXCL16* had higher steady-state expression in secretory cells (supported by MAFF) and was induced in basal, suprabasal and secretory cells by basal/secretory dynamic TFs RREB1 and KLF3. For the basal IDG *CXCL14*, high steady-state, basal-cell expression is maintained by baseline TF MYC, while IFN-increased ETS2 TFA is predicted to downregulate *CXCL14* expression. Ciliated TF ZNF33A is predicted to maintain high baseline expression of *CCL28*, while HESX1 and SP140 upregulate its expression following IFN (panel ii). Secretory-specific, steady-state expression of *CXCL1/2/3/6/8/17* is coordinated by TFs ATF3, KLF3, MESP1, LTF and MAFF (panel iii).

Taken together, our integrated single-cell transcriptomic and chromatin accessibility analysis enabled the construction of GRNs that describe both shared and cell type-associated TF circuits in the HAE at steady-state and following IFNβ stimulation. Our findings provide a genome-scale model for understanding airway homeostasis and offer novel insights into cell type- associated responses that may inform future therapeutic interventions and drug targeting.

## Discussion

Canonical IFN-I signaling and the corresponding gene expression response are central to effective antiviral defense. Across virtually all nucleated cell types, IFN-I engages the same receptor, utilizes shared signaling components, and activates a common set of “primary” TFs (most notably ISGF3 and STAT1:STAT1 homodimers), inducing expression of generally well-defined “shared” ISGs. However, beyond the widely recognized common ISG response, cell type-associated IFN-responsive gene programs have also been observed. Two decades ago, drawing on observations from focused studies, Stark and colleagues proposed that apparent differences in IFN-responsive gene expression and function across cell types could be driven by differential STAT activity, additional signaling pathways (including kinases such as PI3K, p38, ERK and JNK), and complementary non-STAT TFs (including NF-κB, AP-1, IRF1 and others)^93^. Enabled by advances in transcriptomics, particularly single-cell transcriptomics, more recent studies have illustrated the remarkable complexity and diversity of the transcriptional response to IFN (and virus-elicited IFN) across immune cell types, endothelial cells, fibroblasts and epithelial cells^94–100^. scRNA-seq studies of mouse and human airway epithelium (nasal and bronchial) have revealed cell type-associated gene expression programs in response to rhinovirus and SARS-CoV-2 infections^101,102^. While these previous studies have implicated select TFs and/or alternative signaling pathways in contributing to differential IFN-responsive gene expression programs, our study, which integrates single-cell transcriptomic and chromatin accessibility data in context-specific GRN inference, provides a comprehensive view of cell type-associated IFN-responsive gene expression programs.

At single-cell resolution, we characterized the gene expression and chromatin accessibility of human airway epithelium cell types in response to IFNβ and at steady state. These data informed a genome-scale gene regulatory network, which we used to elucidate regulatory mechanisms driving HAE cell type identities and the surprising complexity of their gene expression responses to IFNβ. Accompanying the 548 “shared” interferon-responsive genes, we observed extensive cell type-associated IRGs (nearly 2000 genes), varying greatly in their expression across the HAE cell types. Our results support cell type-associated chromatin priming as an underlying mechanism, as genes induced more strongly in one cell type, for example, are associated with increased steady-state accessibility at the promoters of these genes in that cell type. Previous studies observed that cell type-specific chromatin state drove interferon-stimulated binding of TFs to ISREs^103,104^. Our accessible chromatin analysis supports this, as the cell type-associated regions of IFN-induced chromatin opening were enriched for ISREs, and most of the cell type-associated IFN-opening regions had elevated chromatin accessibility in the relevant cell type at steady state. However, we also identified additional sequence determinants of cell type-associated IFN-induced chromatin opening, including AP-1 motifs that were exclusively enriched in basal and basal-secretory-associated IFN-induced opened chromatin.

We used the GRN to predict regulators of the cell type-associated IRG responses, implicating a general mechanism by which “baseline” TFs collaborate with “dynamic” TFs to orchestrate the differential expression of gene programs across HAE cell types. As expected, the baseline TFs are HAE cell type-associated at steady state, while dynamic TFs can be categorized as shared (e.g., ISGF3, IRFs, STAT1:STAT1, STAT3) and cell type-associated (e.g., POU3F1 in basal cells, ATF3 in secretory cells). The IFN-induced activities of cell type-associated dynamic TFs likely reflect novel crosstalk between well-established IFN signaling components and other intracellular signaling molecules^99^, and, in a tissue context, may also be the product of secondary cell-cell signaling events induced by IFN. While our GRN rediscovers canonical interferon regulators central to the HAE responses observed, it greatly expands the cast of TFs that orchestrate IFN signaling and the initiation of cell type-associated gene expression programs in HAE. Overall, our findings support a model in which cell type-associated IFN responses emerge from the interplay between shared signaling architecture, cell type- and context-dependent TF activities and chromatin accessibility landscapes.

The “HAE IFN regulatory atlas” presented here is expected to be of considerable utility in developing therapeutic strategies for chronic lung diseases, many of which are linked to dysfunction in specific airway epithelial cell types. Moreover, with respiratory virus infection outcomes associated not only with IFN response magnitude but also recently linked to the activity of non-STAT TF responses^101^, these results may inform the design of novel, host-directed antiviral therapeutic strategies.

### Study limitations and future directions

Although there is literature support for many of the predicted regulators of HAE cell type identify and IRGs, the GRN predictions have not been experimentally tested. While TF binding predictions from the ATAC-seq data boost GRN inference accuracy from gene expression data^59,105^, our estimates of “protein TF activity in the nucleus”, used to explain gene expression patterns, are initially based on a combination of TF mRNA abundance and TF binding site prediction from chromatin accessibility data, both imperfect TFA approximations (e.g., due to post-translational modification of TF mRNA products or similarity of DNA sequence recognition by TF family members). For TFA estimation from TFBS data, we initially assumed an activating regulatory interaction between a TF and its TFBS-proximal putative target gene, so there may be inaccuracies in the predicted sign of some regulatory interactions, for predominantly repressor TFs. Thus, measurements of TF activity, in addition to TF perturbation experiments would lead to refinement of the GRN. Our *in vitro* tissue model is limited to epithelial cells (i.e., missing immune cells and immune-epithelial interactions). Thus, the extrapolation of our GRN to *in vivo* HAE tissue needs to be evaluated, and the expansion of the training data to include additional physiological conditions is expected to refine and improve the biomedical relevance of our GRN of the human airway epithelium.

## Acknowledgements

This project was supported by National Institute of Health funding: U01AI150748 (ERM,BRR,CB,LCK,MTW,IM,WZ), R01HL166245 (WZ,ERM), R01AI153442 (ERM), R01AI173314 (ERM,LCK), R01HL151840 (MAS), R01AI151028 (BRR,MAS), R21AI149180 (BRR,MAS), U24HG013078 (MTW), R01AI024717 (MTW,LCK), R01HG010730 (MTW,LCK), R01AR073228 (MTW,LCK), P30AR070549 (LCK,MTW). **Fig. 1A** used BioRender.com. We are also grateful to the directors and teams at Spirovation, affiliated with the Marsico Lung Institute at the University of North Carolina at Chapel Hill, and the MPRI Flow Cytometry and Cell Sorting Facility at the University of Maryland, College Park, for their assistance.

## Data Availability

The data generated in this study have been deposited on the Gene Expression Omnibus (GEO) and will be made available upon manuscript acceptance.

## Supplemental Figure Legends

**Fig. S1.**
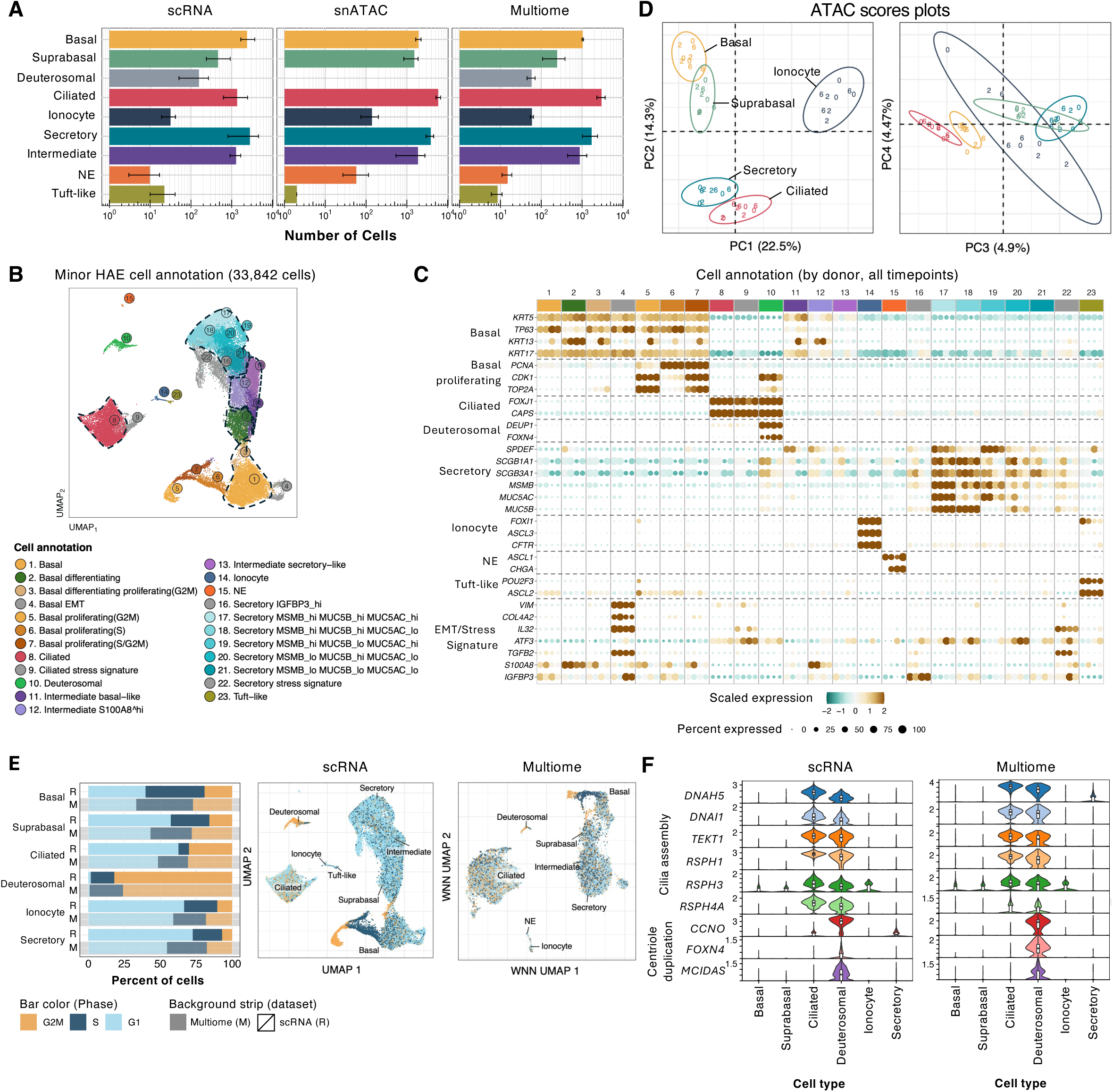
Identification of HAE cell populations at high resolution, related to Fig. 1. **(A)** Average number of cells per population across scRNA-seq, snATAC-seq and sn-multiome-seq datasets. Error bars represent the range across biological donors (n=4 for scRNA, n=3 for snATAC, n=2 for multiome). (**B**) UMAP visualization of and (**C**) gene markers used to identify cell types and subtypes from the scRNA-seq data. Data were normalized by *SCTransform.* (**D**) PCA of pseudobulk (cell type, timepoint, and donor) chromatin accessibility profiles derived from snATAC-seq data. Pseudobulk samples are colored by cell type, with timepoint indicated (“0”, “2”, or “6” hours post-IFN) for each of the three donors. Ellipses represent 95% confidence intervals for each cell type. Axis labels include percent variance explained by each PC in parentheses. (**E**) The percentage of cells in each cell phase (G1, S and G2M) in scRNA-seq and sn-multiome-seq data sets (**Methods**); UMAPs show the cell phase annotations for individual cells. (**F**) Violin plots of normalized gene expression for cilia assembly (*DNAH5, DNAI1, TEKT1M, RSPH1, RSPH3* and *RSPH4A*) and centriole duplication (*CCNO, FOXN4* and *MCIDAS*) genes, as derived from scRNA-seq or sn-multiome-seq data. Data were normalized using *SCTransform*.

**Fig. S2.**
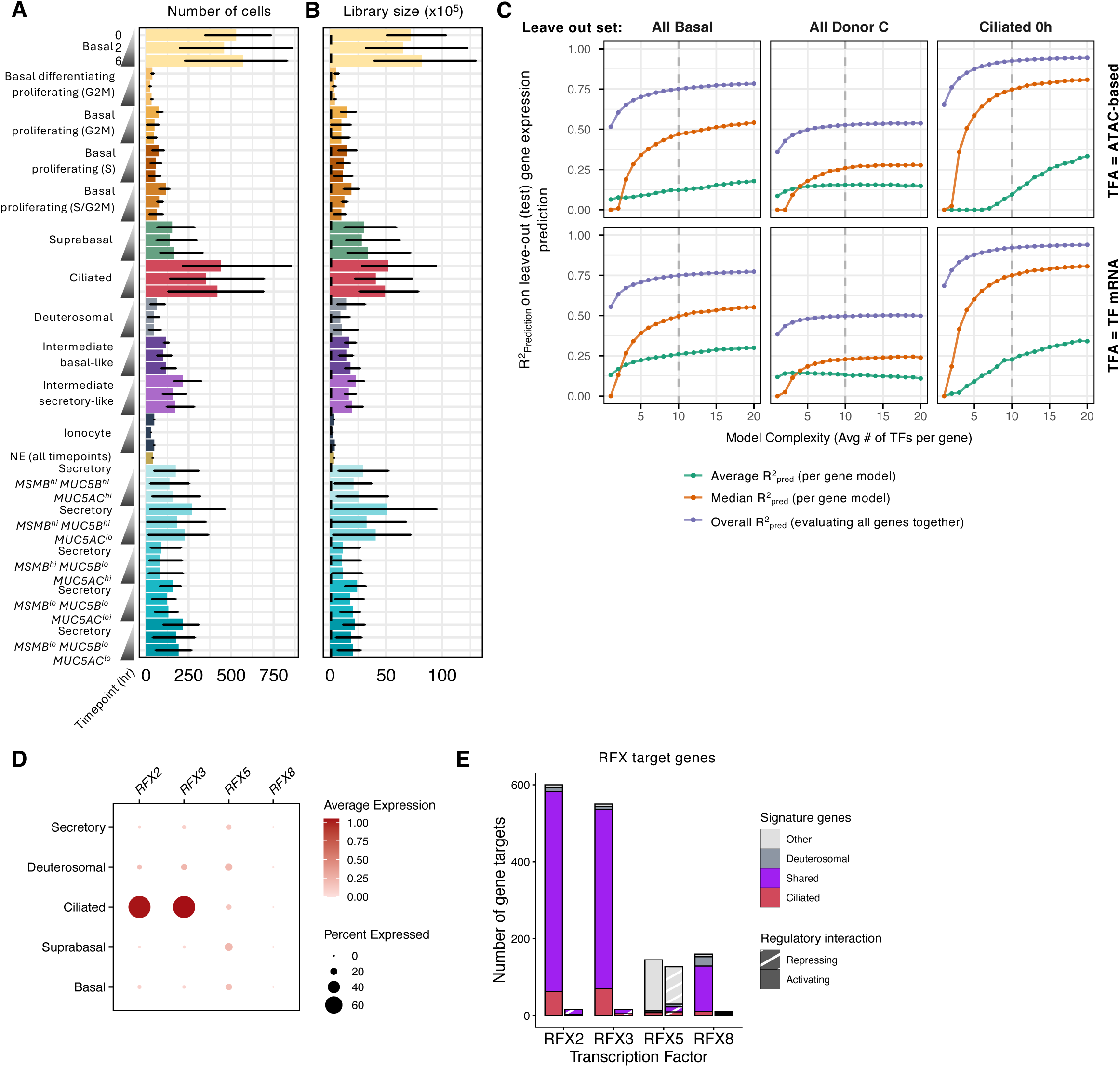
GRN model selection and prioritization of RFX family members, related to Fig. 1 and Fig. 2. The average (**A**) number of cells and (**B**) library size (UMI count) per cell population and timepoint, in the scRNA-seq dataset; error bars represent the range across four donors. (**C**) Out-of-sample gene expression prediction was used to select model complexity (i.e., size or number of TF-gene regulatory interactions in the GRN, reported here as the average number of TF regulators per gene). Our gene expression samples were split into independent sets of training and test samples. Test performance (R^2^ of prediction (**R^2^_pred_**)) was evaluated for three train-test splits of the data, where leave-out test data were (from left to right figure panels): (1) all basal cell pseudobulk samples, (2) all samples from donor C or (3) ciliated pseudobulk samples at steady-state. Across leave-out performance simulations, the plateau in predictive performance (median **R^2^_pred_**) observed at 10 TFs/gene (dashed grey line) justifies the selection of this threshold and the size of the final GRN. (**D**) The expression of *RFX* genes at steady state (normalized using *SCTransform*); dot diameter indicates fraction of cells with transcript detected. (**E**) The number of signature genes (ciliated-unique (red), deuterosomal-unique (grey) or shared (purple)) regulated by each RFX TF; shading indicates activating (solid) or repressing (dashed) regulatory interactions.

**Fig. S3.**
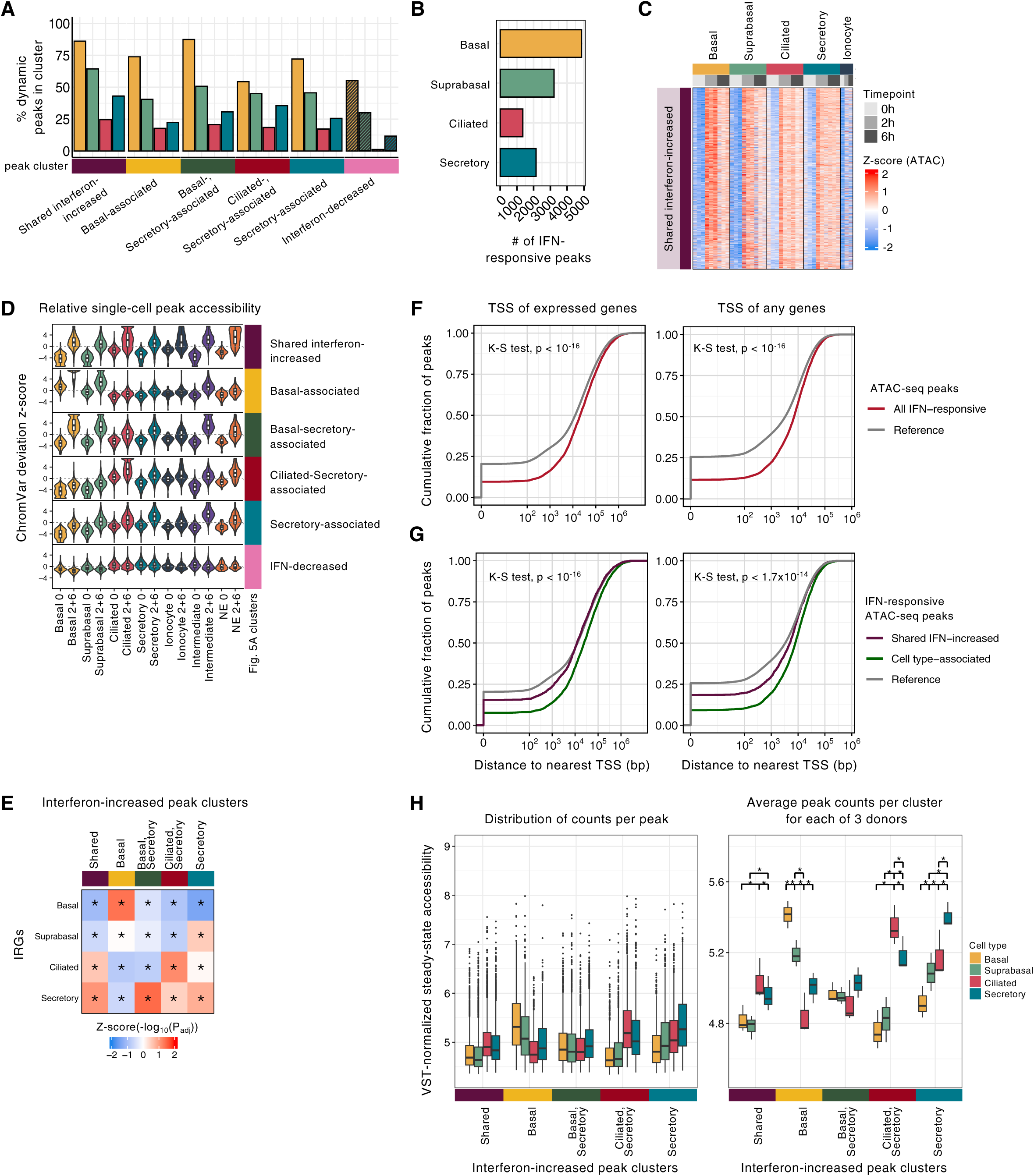
IFN-responsive chromatin accessibility across HAE cell types, proximity of peaks to genes and evidence for steady-state priming, related to Fig. 5. (**A**) The percentage of dynamic peaks in each cluster (from **Fig. 5A**) identified by differential accessible chromatin analysis in each cell type (2 or 6h relative to 0h, P_adj_ < 0.05, |log_2_(FC)| > 1). (**B**) The number of interferon-responsive peaks per cell type, defined as in (**A**). (**C**) Pseudobulk chromatin accessibility of the shared interferon-increased peaks (cluster 1 of **Fig. 5A**), combining n=3 donors per ionocyte timepoint, to boost signal in this rare cell type. (**D**) The accessibility of peak clusters (from **Fig. 5A**) were evaluated in individual cells, using *ChromVar* deviations (z-scores) across rare and abundant cell types at baseline or 2 and 6h. (**E**) Enrichment of peak clusters proximal (±2kb TSS) to IRGs in each cell type (Fisher’s exact test, *P_adj_ < 0.05), z-scored per column. (**F**) The empirical Cumulative Distribution Functions (**eCDFs**) of distances between IFN-responsive or all reference ATAC-seq peaks and the nearest TSS of an expressed gene (left) or any gene (right). (**G**) Similarly, the eCDFs of distances (**G**) between cell type-associated or shared IFN-responsive peaks and nearest TSS of an expressed gene (left) or any gene (right). (**H**) The distribution of steady-state accessibility in basal, suprabasal, ciliated or secretory cells for each interferon-increased peak cluster (left), or averaged per cluster for each donor (right, paired t-test, with Benjamini-Hochberg correction for the four pair-wise comparisons per cluster, *P_adj_ < 0.05).

**Fig. S4.**
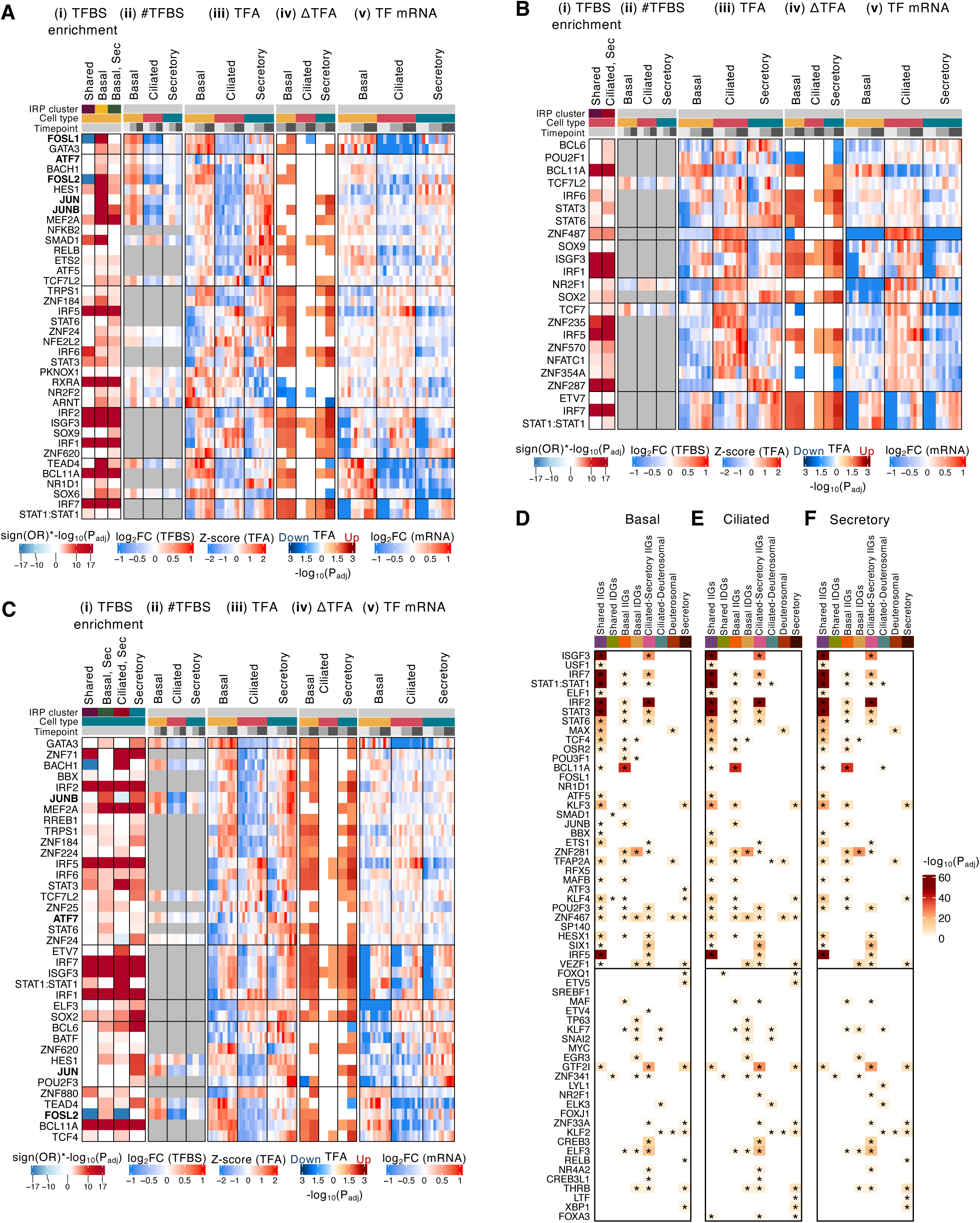
Putative TF regulators of IFN-responsive chromatin and/or chromatin priming proximal to IRGs, related to Fig. 5 and Fig. 6. TFs predicted to regulate IFN-responsive peak clusters in (**A**) basal, (**B**) ciliated and (**C**) secretory cells, showing (i) the enrichment of cell type-specific predicted TFBS (multiplied by the sign of the log_2_OR) in the indicated interferon-response peak (**IRP**) cluster (**Methods**), (ii) the log_2_FC in the number of predicted TFBS relative to the mean, for TFs with a *maxATAC* model available, (iii) the donor-resolved TFA estimates, (iv) the significance of the change in TFA (2 or 6h relative to 0h, based on t-test), multiplied by the sign (increased or decreased TFA), and (v) the log_2_FC in TF mRNA relative to the mean. (**D-F**) Data supporting TFs predicted to prime chromatin proximal to cell type-associated IRGs at steady state. In **Fig. 6B, 6D** panels iii, TFBS enrichment was evaluated using TFBS in the reference ATAC-seq peak set, and these results represent a summation of putative TFBS across cell types. To make statements about priming in particular cell types at baseline, we therefore also evaluated TFBS enrichment using the ATAC-seq peaks detected in each cell type at t=0, so that we could show, for example, that a TF predicted to prime chromatin in basal cells at baseline had chromatin accessibility in its TFBS in basal cells at t=0. Thus, using the t=0 ATAC-seq peaks in (**D**) basal, (**E**) ciliated or (**F**) secretory cells, we depict TFs whose TFBS (from *maxATAC* or motif scanning) are enriched proximally to genes within the IRG cluster in each cell type, respectively; Fisher exact test, *P_adj_ < 0.05; while box shading indicates predicted activator or repressor role, based on TF’s target enrichments in the IIGs or IDGs of each IRG cluster.

**Fig. S5.**
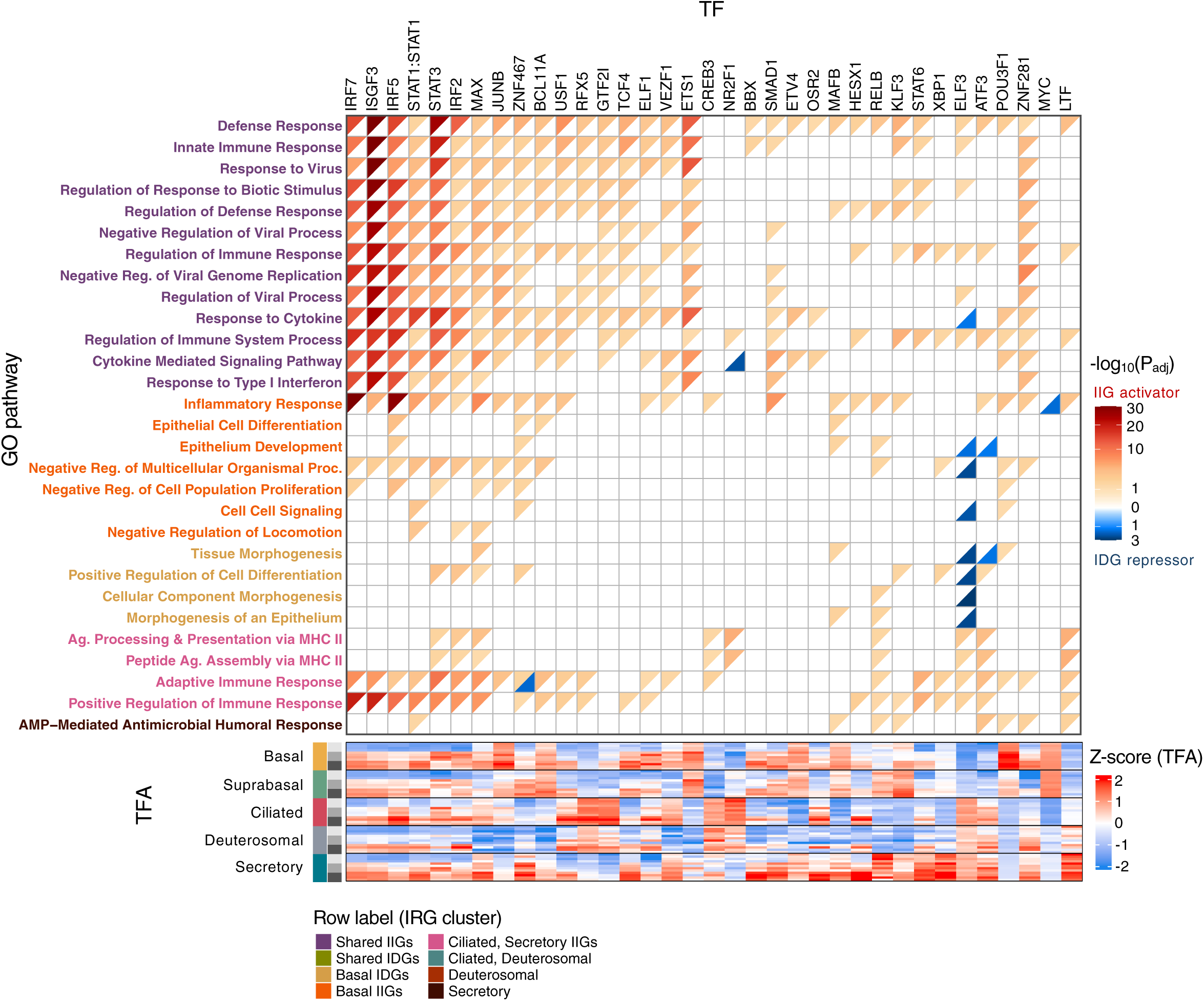
Enrichment of TF targets in IFN-responsive GO terms, related to Fig. 4 and Fig. 6. Enrichment of TF up- (red) or down-regulated (blue) targets in GO biological pathways enriched in IRGs (**Fig. 4J**, Fisher’s exact test, P_adj_ < 0.05). Pathway color indicates IRG cluster. Heatmap showing the donor-resolved TFA estimates across the five queried HAE cell types.

## Methods

### METHOD DETAILS

#### Tissue culture

Normal human tracheobronchial epithelial cells (NHBE, Lonza, cat. # CC-2540S) were used as the cell source for all human airway epithelial (HAE) cultures. Donor IDs “A”, “B”, “C” and “E” were used for single cell genomics experiments, and Donor IDs “A”, “E”, and “H” were included in mucociliary transport assays (available donor metadata included below). NHBE were cultured in submerged phase in Pneumacult^™^-Ex Plus medium (StemCell Technologies, cat. # 05040) before seeding on rat tail collagen type 1-treated permeable Transwell^®^ membrane supports (Corning Inc., 6.5 mm diameter, 0.4 um pore size, cat. # 3470). After reaching confluence, apical medium was removed and cultures were grown at air-liquid interface (ALI) in Pneumacult^™^-ALI maintenance medium (StemCell Technologies, cat. # 05001) as previously described^1061^. All HAE cultures were maintained at ALI for >4-weeks to ensure proper differentiation to polarized, pseudostratified epithelium with motile cilia and mucus secretion. All cell cultures were maintained at 37C with 5% CO_2_. HAE cultures from “donor B” were evaluated in duplicate independent differentiations.

| Code | Age | Sex | Race | Alcohol | Smoking |
| --- | --- | --- | --- | --- | --- |
| Donor A | 65 | M | Caucasian | N | Y |
| Donor B | 73 | F | Caucasian | N | N |
| Donor C | 66 | F | Black and Hispanic | N | N |
| Donor E | 52 | M | Hispanic | Y | N |
| Donor H | 55 | F | Hispanic | N | N |

#### IFN stimulation and dissociation of HAE cultures

Prior to IFNβ stimulation, the apical surfaces of all HAE cultures were washed with 200 μL Dulbecco’s PBS (no Ca2+, no Mg2+, Gibco cat. # 14190250) at 37C for 10m. HAE cultures were stimulated with IFNβ by replacing the basolateral medium with Pneumacult^™^-ALI maintenance medium supplemented with 1 nM recombinant human interferon beta 1a (PBL Biosciences, cat. # 11415-1) at 2h and 6h prior to harvest. For the 0h timepoint, HAE cultures were maintained in Pneumacult^™^-ALI maintenance medium without IFNβ supplementation.

To initiate dissociation, HAE cultures were transferred to TrypLE dissociation solution (Gibco, cat. # 12-604-013) with 10 μM ROCK (Rho-kinase) inhibitor Y-27632 (Enzo Life Sciences, cat. # ALX-270-333-M025), adding 500 μl to the basal chamber and 100 μl to apical surface. Following incubation for 5m at 37C, the apical solution was aspirated and quenched in a pre-warmed trypsin neutralizing solution (Lonza, cat. # CC-5002) with ROCK inhibitor Y-27632 (10 μM final concentration). Dissociation proceeded in sequential rounds of 1.5m at 37C, pooling harvested fractions until efficient dissociation was confirmed by microscopy. For scRNA and snATAC assays, cells were pelleted at 250 x g for 5m at RT and resuspended in 500 μl Dulbecco’s PBS with 10 uM Y27632 ROCK inhibitor cocktail (DPBS + RI). Cell count and viability were assessed using trypan blue dye exclusion on a Countess II Automated Cell Counter (Invitrogen) before proceeding to respective downstream applications.

#### Single cell library preparation and sequencing

##### scRNA-seq

For each sample, cell suspensions were first labelled with barcoded MULTI-seq oligos as described^107^. Briefly, 250,000 cells per sample were resuspended in 180 uL of DPBS + RI and incubated with 20 ul prepared lipid modified oligo (**LMO**) barcode for 5m on ice. Following addition of MULTI-seq co-anchor solution and subsequent incubation for 5m on ice, the cell suspension was washed 3 times in DPBS + RI with 1% w/v BSA and centrifuged at 250 x g for 3m at 4C. Cells were then filtered through a 10 um filter (PluriSelect, cat. #431001050) into a clean 1.7 mL microcentrifuge tube. Cell count and viability were re-assessed using trypan blue dye exclusion on a Countess II Automated Cell Counter (Invitrogen), requiring an estimated viability of above 80% to proceed. MULTI-seq barcoded cell pools were combined and adjusted to balance the representation of each sample. 10X Genomics 5’ libraries were prepared using the Chromium Next GEM Single Cell V(D)J Reagent kit v1.1, with modifications to isolate the MULTI-seq barcodes as described^107^ in McGinnis et al. 2019. 10 PCR cycles were used to amplify MULTI-seq barcode libraries from 3.5 ug input cDNA. Gene expression and MULTI-seq libraries were sequenced separately for series 1 data (donors A, B, and C) and together for series 2 data (donors B and E). Libraries were sequenced on either an Illumina NextSeq 550 (series 1 data) or Illumina NovaSeq 6000 (series 2 data) instrument with the following read configurations. Series 1 - read 1: 8 cycles, read 2: 36 cycles, index read 1: 8 cycles, index read 2: 0 cycles. Series 2 - read 1: 29 cycles, read 2: 92 cycles, index read 1: 10 cycles, index read 2: 0 cycles.

##### snATAC-seq and snMultiome-seq

For snATAC-seq, 200,000 cells were transferred to a sterile 1.7 mL microcentrifuge tube and diluted to a final concentration of 1M cells / mL. The cell suspension was supplemented with bovine serum albumin stock solution (Miltenyi, cat. # 130-091-376) at a final concentration of 0.04% w/v. Library preparation for snATAC-seq proceeded as per the 10X Genomics Chromium Nuclei Isolation Kit User Guide and Chromium Next GEM Single Cell ATAC Reagent Kits User Guide (v2, revision A). Resulting libraries were sequenced on an Illumina NovaSeq 6000 instrument with the following read configuration: read 1: 51 cycles, read2: 51 cycles, index read 1: 8 cycles, index read 2: 24 cycles. For a subset of samples, joint gene expression and chromatin accessibility profiling was performed using snMultiome-seq. Cells were isolated and resuspended in DBPS with ROCK inhibitor as in the snATAC-seq protocol. Library preparation then proceeded as described in the Chromium Next GEM Single Cell Multiome ATAC + Gene Expression User Guide for joint gene expression and chromatin accessibility profiling. Gene expression libraries were sequenced on an Illumina NovaSeq 6000 instrument with the following read configuration: read 1: 29 cycles, read 2: 91 cycles, index read 1: 10 cycles, index read 2: 10 cycles. ATAC libraries were sequenced on an Illumina NovaSeq 6000 instrument with the following read configuration: read 1: 51 cycles, read 2: 51 cycles, index read 1: 8 cycles, index read 2: 24 cycles.

#### RNA FISH

HAE cultures were stimulated with IFNβ, as above, for 6h prior to fixation in 4% paraformaldehyde. HAE cultures were then embedded in paraffin and 5-micron thick sections were prepared on slides by Spirovation (associated with the Marsico Lung Institute at the University of North Carolina at Chapel Hill). Fluorescent in situ hybridization for the simultaneous detection of *MX1* and *CXCL*10 mRNA was performed using the RNAscope Multiplex Fluorescent Reagent kit V2 (Advanced Cell Diagnostics, cat. # 323110). Briefly, slides were baked (60C; 1 h), deparaffinized with xylene, and then washed with 100% ethanol and air-dried (60C; 5m). Slides were next treated with RNAscope Hydrogen Peroxide solution at room temperature for 10m prior to target retrieval as per the manufacturer’s instructions. HAE culture sections were then covered with RNAscope Protease Plus and incubated at 40C for 15m, after which they were rinsed in water, and incubated with RNAscope probes (Hs-MX1-C1, cat. #403831; Hs-CXCL10-C3, cat. #311851-C3) at 40C for 2h. After washing, a series of amplification steps were performed at 40C (AMP1 and AMP2 for 30m each; AMP3 for 15m) with two washes in between each amplification. HAE sections were then treated with HRP-C1 for 15m at 40C, washed twice, and incubated with TSA plus Fluorescein/Cy3 fluorophore (1:1500, Perkin Elmer cat. # NEL760001KT) for 30m to visualize the *MX1* mRNA (Channel 1 at 546 nm). Following two washes, a 15-min incubation with RNAscope HRP blocker, and two more washes, the sections were treated as above, this time using HRP-C3 and TSA plus Fluorescein/Cy5 fluorophore (1:1500, Perkin Elmer cat. #NEL760001KT) to visualize the *CXCL10* mRNA (Channel 3 at 647 nm). A final 15m incubation with RNA HRP blocker and series of wash steps were then performed, and sections treated with DAPI for 30s, prior to being cover-slipped with ProLong Gold Antifade Mountant. Images were acquired with a Zeiss Axio Observer 3 inverted fluorescence microscope equipped with a Zeiss AxioCam 503 mono camera and Zen 2.3 Lite imaging software.

#### Flow cytometry

HAE cultures were stimulated with 1 nM IFNβ on both the apical and basolateral surfaces for 24h and 48h before being dissociated to a single cell suspension as described above (see “IFN Stimulation and Dissociation of HAE cultures”). Isolated HAE cells were passed through a 10-micron filter (PluriSelect, cat. #431001050) and then resuspended in BD Cytofix/Cytoperm solution (BD Biosciences, cat. # 554714). Staining with anti-acetylated alpha tubulin-AlexaFluor 647 (1:100; Santa Cruz cat. # sc23950-AF647) and anti-CD38-PE (5 μL / test; Invitrogen cat. # 12-0389-42) was performed as per the BD Cytofix/Cytoperm kit instructions. Cells were then pelleted at 500 x g for 3m and resuspended in flow cytometry running buffer (0.1% BSA, 10 mM HEPES, PBS). All data were acquired in the MPRI Flow Cytometry core facility at the University of Maryland using a FACSCanto II and processed with FACSDiva and FlowJo software v10.10.0.

#### Mucociliary transport (MCT) measurements

HAE cultures were washed with PBS one week prior to MCT analysis to ensure the establishment of a robust mucus barrier. MCT was then assessed using fluorescence video microscopy with single particle tracking. At the 0h timepoint, carboxylate-modified fluorospheres (FluoSpheres^TM^, 2.0 mm, red 580/605; Invitrogen cat. # F8826) were diluted 1:3000 in DPBS, and 20 µl were added to the apical surface of each culture. Cultures were then incubated at 37C for 15m to allow the beads to settle atop the mucus gel prior to imaging. Fluorescence video microscopy was performed at 10X magnification on a Zeiss LSM 800 equipped with an Axiocam 702 mono camera and Zen 2.3 imaging software blue edition (widefield mode; 20 s videos; 100ms exposure time). MCT video capture was repeated to collect input from a total of 3-4 different, and randomly selected regions of the culture. After imaging at the 0h timepoint, cultures were maintained with or without IFNβ (2 nM; PBL Assay Science, Cat #: 11415-1) in the basolateral compartment, along with CD38 inhibitor (78C; 1 mM; TOCRIS Bioscience, cat #:6391) or dimethylsulfoxide (DMSO (vehicle control); 1 mM; ATCC, cat. # 4-X) in both the apical and basolateral compartments. For apical treatments, reagents (78C; DMSO) were diluted in DPBS and 10 μL was applied to the culture surface. After 24h at 37C, additional beads were added to the apical surface and MCT videos were taken, as above. Analysis of MCT videos was performed using MATLAB, as previously described^93^ and resulting data were graphed using GraphPad Prism Software v10.4.2.

### QUANTIFICATION AND STATISTICAL ANALYSIS

In figures, asterisks denote significance based on P_adj_ < 0.05, as determined by appropriate statistical testing (e.g., two-tailed t-tests for continuous variables, negative binomial distribution for genomic counts data), as indicated in the Figure Legends and detailed below. The Benjamini-Hochberg procedure was used to correct for multiple hypothesis testing.

#### Single-cell RNA-seq processing

##### Alignment, quality control, and doublet removal

Reads were aligned to the 10x Genomics GRCh38 reference genome, filtered and UMIs were counted using Cell Ranger Multi v5.0.0^108^. Cell Ranger outputs and downstream single-cell analyses were processed and performed with Seurat v4.1.1^109^ and v5^110^ for all sequencing modalities. High-quality cells were retained if they had (1) at least 5,500 unique transcripts (UMI) following demultiplexing and (2) the percentage of mitochondrial reads was less than 20%. Next, we filtered out contaminating doublets. For scRNA-seq samples, we removed doublets detected during sample demultiplexing based on MULTI-seq oligos. We also applied *scDblFinder* v1.18.0^111^ to identify additional intra-sample doublets (using default parameters without artificial doublet generation).

##### Sample integration and clustering

scRNA-seq data were normalized using *SCTransform* v2^109,112^ and integrated using a reciprocal PCA strategy implemented in Seurat v4.1.1. Integration was performed across donor, timepoint and experimental batch factors using 4000 highly variable features and was based on the RPCA workflow in the Seurat documentation. Louvain clustering with a resolution parameter of 2.2 was used for the first round of clustering and per-cluster QC. At this clustering resolution, 5 low-quality clusters were excluded based on low UMI counts, high % mitochondrial content, or suspected doublet status, leaving 34 remaining clusters. At this point, data were re-clustered using the Louvain algorithm, at a range of resolutions (0.2 to 3) to assess cluster stability. Cell type annotation was performed in a semi-supervised manner in two stages. First, major HAE populations were identified using label transfer from scRNA-seq of HAE cultures^48^. All major cell types exhibited stable clustering at a resolution of 1.4, with the exception of ciliated cells, which were annotated using a resolution of 1.0. For each major group, subclusters were individually assessed based on marker gene expression and assigned a minor group label (**Fig. S1C**); some highly similar subclusters were merged. Then, for candidate tuft-like cells and ionocytes, subclustering was performed (again with the Louvain algorithm and a resolution of 1.4), Putative tuft-like cells were distinguished from ionocytes based on expression patterns of the established tuft-like regulator *POU2F3* and ionocyte markers *ASCL3* and *CFTR*. The final annotated dataset contains 23 minor group labels and 33,842 cells (**Fig. S1B-C**). For all downstream analyses, we excluded several minor clusters that showed signatures of stress, epithelial-mesenchymal transition, or high *IGFP3* expression (clusters 4, 9, 12, 16 and 22, **Fig. S1C**).

#### Single-nuclei ATAC-seq processing

##### Alignment, quality control and doublet removal

Initial alignment, barcode filtering and transposase cut site identification were performed using Cell Ranger ATAC v2.0.0^113^ using 10x Genomics GRCh38 reference genome. Following the *Signac* workflow^114^, we kept high quality barcodes if they had a minimum TSS enrichment score of 2, percent fragments in blacklist region less than 3%, a maximum nucleosome signal score of 2, percent reads in peaks greater than 40%, and number of ATAC fragments ranging from 2,000 to 200,000 per cell (nCount), and at least 1,000 cut sites in the initial peak set (nFeature). Next, we identified potential doublets, using fragment files as input to *ArchR* v1.2.0^115^. Expected doublet rates were obtained from the 10x Genomics website (Chromium Next GEM Single Cell ATAC Reagent Kits User Guide (v2, revision A)). The estimated number of doublets was calculated by multiplying the expected doublet rate with the number of recovered cell barcodes in that sample. Nuclei were ordered based on their predicted doublet score, and the expected number of doublets were removed from the highest-scoring nuclei.

##### Sample integration, clustering, and identification of reference peak set

To integrate snATAC-seq samples, we first defined a common reference peak set by running *MACS2* v2.1.4^116^ on our fragment cut-site data. To generate cut sites, we split each fragment into its two transposase insertion points, merged them into a BED file, then excluded ENCODE blacklist regions^117^. For each time point and replicate, we smoothed cut sites by extending ±25bp and called peaks with *MACS2* using ‘--shift -25 --extsize 50 --q 0.001 --keep-dup all’ parameters, effectively generating 50bp fragments centered on each cut site. We then generated a peak-by-cell count matrix, applied TF-IDF normalization (*Seurat* ‘*RunTFIDF*’), and reduced dimensionality via latent semantic indexing (**LSI**) (*Seurat* ‘*RunSVD*’), retaining the top 50 singular vectors. Finally, we dropped the first LSI component, due to per-cell correlation with a technical covariate: sequencing depth (Pearson correlation > 0.8. Then, we merged the samples and applied *Seurat*’s anchor-based integration to the LSI embeddings (*Seurat* ‘*IntegrateEmbeddings*’).

Following integration, we performed label transfer to nuclei in the snATAC-seq dataset. We counted the number of cut sites (using the +/-25bp extension) overlapping gene promoters using *Seurat*’s ‘*GeneActivity*’ function (2,000bp upstream to 500bp downstream of the TSS). We log-normalized the counts and used PCA for dimensionality reduction. Labels were then transferred using ‘*FindTransferAnchors*’ and ‘*TransferData*’, and nuclei were annotated based on their highest prediction score. For each cluster, we labelled nuclei based on the most frequent cell type in that cluster.

After identifying major and minor populations, we called peaks for each cell type and timepoint, aggregating signal from all donors, using the same parameters described above. The resulting peaks were combined into a final reference peak set, to generate a peak-by-cell count matrix used in downstream analyses. Across cell types and time points, this pipeline yielded 459,941 peaks from the snATAC-seq dataset.

##### Visualization of chromatin accessibility signal tracks

For visualization of chromatin accessibility tracks, signal tracks were normalized per million reads using the SPMR flag (*--SPMR*) from *MACS2*, which outputs normalized bedGraphs. Normalized bedGraphs were then converted to BigWig format using the *bedGraphToBigWig* function from *UCSC tools* v3.8.0^118^. All chromatin track data was visualized using *IGV* v2.16.0^119^ and group auto-scaled across tracks.

#### Single-nuclei multiome-seq processing

Alignment, quality control and doublet removal were performed on both gene expression (RNA) and chromatin accessibility (ATAC) assays, as described above. Cell Ranger ARC v2.0.0 was used for alignment and initial peak calling. High-quality barcodes were identified from both assays according to the following: for the ATAC assay, we retained nuclei with TSS enrichment score greater than 2, percent fragments in blacklist region less than 3%, percent reads in peak regions greater than 40%, nucleosome signal score less than 2, at least 1,000 cut sites in the initial peak sets (nFeature), and number of ATAC fragments ranging from 1,000 to 100,000. For the gene expression assay, we retained nuclei with number of UMIs ranging from 2,000 to 25,000 and percent mitochondrial reads less than 50%. To identify and remove potential doublets, we took an ensemble approach and ran *ArchR* v1.2.0 and *scDblFinder* v1.18.0 on the ATAC and RNA assays, respectively. Nuclei were ranked according to the doublet confidence for each of the two methods. Mirroring snATAC-seq analysis, expected doublet rates were obtained from the 10x Genomics website (Chromium Next GEM Single Cell Multiome ATAC + Gene Expression User Guide). The expected number of doublets was calculated by multiplying the expected doublet rate with the number of recovered cell barcodes in that sample, and the expected number of doublets were removed based on rank of doublet scores for each assay.

Next, we leveraged both ATAC and RNA assays to cluster nuclei. First, samples were integrated per assay using *Harmony* v1.2.1^120^. For the ATAC assay, we performed TF-IDF and LSI, then, using dimensions 2 through 30, as input to *Harmony*. For the RNA assay, we performed PCA on the adjusted, normalized data (*ScaleData* from Seurat), and used PCs 1-30 as input to *Harmony*. For each assay, we treated each experiment as a “batch” variable (*RunHarmony*). Next, we used *Seurat* to construct a weighted nearest neighbor (**WNN**) graph using the ATAC- and RNA-harmonized assays, which informed clustering and UMAP calculation. We identified major and minor cell types in each sample using *Seurat*’s anchor-based label transfer from the scRNA-seq gene expression data. Finally, we identified a multiome reference peak set of 197,624 peaks, following the procedure described for snATAC-seq data above.

#### Pseudobulk analysis of gene expression data

Pseudobulk gene expression samples were generated by summing raw counts from cells, aggregated per cell type, donor and time point. The pseudobulk gene expression matrix was limited to genes detected in at least 5% of cells pertaining to at least one cell type-time point combination.

For PCA and differential gene expression analysis, we required a minimum pseudobulk library size of 100k transcripts; thus, rare cell types (e.g., NE and Tuft-like), were excluded from these analyses. We considered six cell types (basal, suprabasal, ciliated, deuterosomal, ionocyte, secretory). “Secretory” were a combination of clusters 17-21, while basal cells were limited to cluster 1 and excluded proliferating clusters (3-7) (**Fig. S1C, G**). From scRNA-seq, the final dimensions of the resulting gene expression matrix was 15,881 genes by 72 pseudobulk samples (n = 4 donors, 6 cell types, 3 timepoints) (**Table S1B**). Raw counts served as inputs to differential expression analyses. For data visualization, the pseudobulk gene expression data (raw counts) were normalized using the variance-stabilizing transformation (**VST**) from the *DESeq2* v1.44.0^121^, setting “*blind*=TRUE”. The resulting matrix was then batch-corrected using ComBat from the *sva* v3.52.0^122^, with donor as the batch covariate.

For GRN inference from the scRNA-seq data, we leveraged heterogeneity within larger populations and combined time points per donor for rare NE cells to achieve the minimum library size (**Fig. S2A-B**). Prior to combination of timepoints for NE cells, the full gene expression matrix was batch-corrected using Combat-seq^123^ from *sva* v3.52.0, then NE timepoints were summed per donor. The resulting gene expression matrix (196 pseudobulk conditions across 17 cell populations, 3 timepoints and 4 donors) was VST-normalized as described for data visualization (**Table S1A**).

##### Principal Component Analysis

We performed PCA on the z-scored, batch-corrected VST gene expression counts matrices from scRNA-seq (**Fig. 1B**). Annotation of genes in the loadings plots were based on selection from the literature and unbiased approaches. For unbiased annotation, we labeled the top-3 genes with greatest absolute contribution to each PC, in addition, we labeled the top-3 genes with the largest pairwise contributions to PC1-2 (correlating with basal/suprabasal cell samples) or PC3-4 (correlating with IFN dynamics). From the literature, we added markers of basal and suprabasal (*TP63*, *KRT14* and *KRT13*), ciliated (*FOXJ1* and *RFX3*), deuterosomal (*FOXN4* and *DEUP1*), secretory (*SPDEF*, *MUC5AC* and *MUC5B*) and ionocytes (*FOXI1* and *CFTR*) to PC1-2 loadings, and canonical ISGs *ISG15*, *ISG20*, *OAS1* and *IDO1* to PC3-4 loadings.

##### Identification of interferon responsive genes (IRGs)

We used *EdgeR* v3.40.2^124^, with the one-factor design “*gene* ∼ *celltype_timepoint*”, to identify IRGs that were differential between 0h and the 2 or 6h post-IFN stimulation, in at least one of the following cell types (basal, suprabasal, ciliated, secretory, deuterosomal cells or ionocytes), using stringent criteria: FDR = 5%, |log_2_FC| > 1 (n=4 donors). Gene expression log_2_-fold changes were estimated using *DESeq2*’s *ashr*^125^ Bayesian shrinkage (**Table S5A**).

##### Definition of signature genes in each cell type

To identify steady-state gene signatures for each cell type (e.g., **Fig. 1C**), we applied criteria^40,61^ designed to identify upregulated and downregulated genes defining highly related cell types at steady state, allowing for partially overlapping gene signatures among cell types. We used *DESeq2*, with the one-factor design “*gene* ∼ *celltype_timepoint*” and the permissive differential analysis criteria: |log_2_FC| > 0.58, FDR = 10%. Genes were limited to those detected in at least 5% of cells. A cell type’s “upregulated signature genes” consist of genes (1) upregulated in that cell type relative to at least one other cell type and (2) not downregulated in that cell type relative to any other cell type, at t=0h. We similarly defined downregulated signature genes. Gene expression log_2_FC were estimated using *DESeq2*’s *ashr*^125^ Bayesian shrinkage (**Table S1A**).

#### Pseudobulk analysis of chromatin accessibility data

For differential analysis, we required a minimum pseudobulk library size of 1M reads, similarly excluding rare cell types (e.g., NE, tuft-like) whose library sizes did not reach the threshold after aggregating across timepoints and/or donors. For ionocytes, timepoints were aggregated per donor to reach the target library size and treated as t=0. Thus, from the snATAC-seq dataset, the final dimensions of the accessibility matrix were 459,941 peaks by 39 pseudobulk samples (n = 3 donors, 5 cell types, 3 timepoints or (for ionocytes) one aggregated timepoint). We additionally generated an snATAC-seq pseudobulk accessibility matrix without aggregating donors in ionocytes for data visualization and PCA to assess reproducibility (**Fig. 1D**, **S1D**).

To leverage the deuterosomal cell accessibility from sn-multiome-seq, we retained deuterosomal cell and ionocyte conditions despite the smaller library sizes, and combined timepoints per donor for all cell types, to assess reproducibility across donors. Thus, the multiome accessibility matrix dimensions were 197,669 peaks by 12 pseudobulk samples (n = 2 donors, 6 cell types) (**Table S2A-B**).

Raw counts served as inputs to differential accessibility analyses.

For data visualization, the pseudobulk accessibility matrices of both datasets (raw counts for snATAC-seq but counts down-sampled to 392,513 fragments for sn-multiome-seq representing the smallest pseudobulk library) were normalized using the variance-stabilizing transformation (**VST**) from the *DESeq2* v1.44.0^121^, setting “*blind*=TRUE”. These matrices were subsequently batch-corrected using ComBat following DESeq2 VST normalization. We used downsampling for sn-multiome-seq data visualization to ensure that the technical covariate (library size) did not drive clustering of cell types in the PCA (**Table S2B**).

##### Principal Component Analysis

We performed PCA on the z-scored, batch-corrected VST chromatin accessibility counts matrices from snATAC-seq (**Fig. S1D**) and sn-multiome-seq (**Fig. 1I**).

##### Identification of interferon responsive accessible chromatin regions (“peaks”)

We used *EdgeR* v3.40.2^124^, with the one-factor design “*accessibility* ∼ *celltype_timepoint*”, to identify peaks that were differential between 0h and 2 or 6h post-IFN stimulation, in at least one of the following cell types (basal, suprabasal, ciliated or secretory cells), using stringent criteria: FDR = 5%, |log_2_FC| > 1 (n=3 donors) (**Table S6A**).

##### Definition of signature peaks in each cell type

Mirroring gene expression, we identified steady-state peak signatures for each cell type (e.g., **Fig. 1C**), allowing for partially overlapping peak signatures among cell types (**Table S6A**). To identify signature peaks in ionocytes, we aggregated counts across timepoints for each donor. We used *DESeq2*, with the one-factor design “*accessibility* ∼ *celltype_timepoint*” and the permissive differential analysis criteria: |log_2_FC| > 0.58, FDR = 10%. A cell type’s “signature peaks” consist of peaks (1) with elevated accessibility in that cell type relative to at least one other cell type and (2) without lower accessibility in that cell type relative to any other cell type, at t=0h. We similarly defined signature peaks with lower accessibility. We identified peak signatures for basal, suprabasal, ciliated, secretory cells and ionocytes from the snATAC-seq dataset (**Fig. 1C**) and expanded this selection to include deuterosomal cells for the multiome-seq dataset (**Fig. 1G**).

##### Prediction of TFBS using maxATAC neural network models

We generated *in silico* TFBS predictions with *maxATAC* v1.0.6^44^ for a total of 27 cell type-timepoint conditions (donor-resolved [n = 3] basal, ciliated, secretory cells at each timepoint: 0, 2, 6h), where pseudobulk library sizes achieved the recommended >5M cut sites. Of the 127 human TFs with *maxATAC* models available, 105 TFs were nominally expressed in HAE and we simulated genome-wide TFBS predictions for these models: ARID3A, ARNT, ATF2, ATF3, ATF7, BACH1, BHLHE40, CEBPB, CEBPD, CEBPZ, CREB1, CREM, CTCF, E2F8, E4F1, EGR1, ELF1, ELF4, ELK1, ESRRA, ETS1, ETV6, FOS, FOSL1, FOSL2, FOXA1, FOXK2, FOXM1, FOXP1, GATA3, GATAD2B, HES1, HSF1, IRF3, JUN, JUNB, JUND, KLF5, LEF1, MAFF, MAFK, MAX, MAZ, MBD2, MEF2A, MNT, MXI1, MYB, MYBL2, MYC, NFATC3, NFE2L2, NFIA, NFIC, NFXL1, NFYA, NFYB, NKRF, NR2C1, NR2F1, NR2F2, NR2F6, NRF1, PAX8, PBX3, PKNOX1, RELA, REST, RFX1, RFX5, RUNX1, RXRA, SIX5, SKIL, SMAD1, SMAD5, SOX6, SP1, SREBF1, SREBF2, SRF, STAT5A, TBP, TCF12, TCF7, TCF7L2, TEAD4, USF1, USF2, YBX1, ZBED1, ZBTB33, ZBTB40, ZBTB7A, ZFX, ZHX2, ZKSCAN1, ZNF143, ZNF217, ZNF24, ZNF274, ZNF282, ZNF569, ZNF592, ZNF687.

##### Prediction of TFBS using TF motif scanning

Using our reference peak set derived from the snATAC-seq or sn-multiome-seq datasets, we scanned sequences within peaks for occurrences of TF motifs, using the CIS-BP motif collection v2.00^126^ and FIMO^127^ (Meme suite v5.4.1), with a raw p-value threshold of 1E-5 and a first-order Markov background model.

#### Gene regulatory network inference

##### Selection of target genes

Using the VST-normalized, batch-corrected pseudobulk gene expression matrix (described above), we built models for 12,417 genes, which were differentially expressed in at least one comparison between cell types at a particular timepoint (e.g., ciliated cells at 2h versus basal cells at 2h), or across timepoints within a particular cell type (e.g., ciliated cells at 2h versus steady state, *DESeq2*, |log_2_FC| > 0.58, FDR = 10%, **Table S3A**).

##### Selection of potential regulators

From a list of 1,639 putative human TFs^128^, we limited potential regulators to the union of TFs that exhibit high variability in their expression (coefficient of variation > 40%) or appear as a signature TF in any condition. The resulting list of potential regulators consisted of 722 TFs. In addition, we considered two TF complexes as potential regulators: ISGF3 and the STAT1:STAT1 homodimer. For ISGF3, we used a representative ISRE motif (MA05517.1 from JASPAR^129^) and, for STAT1:STAT1 homodimers, representative GAS motifs (MA0137.2 and MA0137.3 from JASPAR). For GRN models constructing using TF mRNA as an estimate of TFA (see below), ISGF3 complex activity was approximated as the minimum of *STAT1*, *STAT2* and *IRF9* expression^130^. We removed CTCF as a potential regulator, due to its role as an insulator.

##### Prior network construction

We derived a “prior” network of TF-target gene interactions from the snATAC-seq data^40,131^. Using *maxATAC* TFBS predictions (when available for a TF) or TF motif scanning in accessible chromatin (otherwise), TFs were linked to target genes if a predicted TFBS was detected within ±10kb of the gene body. Interaction weights for each TF were Frobenius-normalized.

##### GRN inference with the Inferelator

We model gene expression as a multivariate linear combination of TFAs:

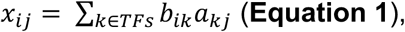

such that 𝑥*_ij_* is the expression of gene *i* in pseudobulk *j*, 𝑎*_kj_* is the TFA of TF *k* in pseudobulk *j*, and 𝑏*_ik_* is the coefficient, learned by the model, which represents the regulatory effect of TF *k* on gene *i* ^59,130^. Ideally, TFA would correspond to a measurement of nuclear protein TF activity, but our experimental design (scRNA/ATAC-seq) did not directly measure protein TFAs (𝑎_*kj*_). Thus, we estimate TFA using two complementary approaches, the combination of which boosts GRN accuracy^59^:

(1) TF mRNA as proxy for TF activities, a reasonable assumption for TFs whose main mode of regulation is transcriptional,

(2) prior-based TFA, estimated using:

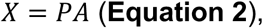

where 𝑋 ∈ ℝ^|genes|×|samples|^ is the gene expression matrix, 𝑃 ∈ ℝ^|genes|×|TFs|^ is the prior network, and 𝐴 ∈ ℝ^|TF’|×|samples|^ is the TFA matrix obtained from solving **Equation 2**^132^. We first generate two GRNs based on either TF mRNA or prior-based TFA, learning coefficients 𝑏*_ik_* by optimizing:

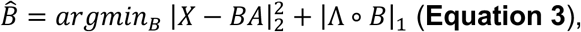

such that 𝑋 ∈
ℝ^|genes|×|samples|^ is the gene expression matrix, 𝐴 ∈ ℝ^|TF’|×|samples|^ is the TFA matrix, 𝐵 ∈ ℝ^|genes|×|*TFS*|^ is the matrix of TF-gene regulatory interaction coefficients, and Λ ∈ ℝ^|genes|×|*TFS*|^ is the matrix of nonnegative penalties associated with each interaction^59^. The first term in **Equation 3** corresponds to model fit, while the second term is an adaptive LASSO regularization term, to incorporate the assumption that the GRN is sparse (only a small subset of TFs regulate a particular gene) and “prior” knowledge (TFBS predictions from snATAC-seq data) to guide inference. As we seek to minimize **Equation 3**, a smaller penalty Λ_*i,k*_ is used for TF-gene interactions supported by the ATAC-prior. Specifically, Λ_*i,k*_ = 𝑏𝑖𝑎𝑠 ∗ 𝜆, where 𝑏𝑖𝑎𝑠 = 0.5, for TF-gene interactions supported by the prior and 𝑏𝑖𝑎𝑠 = 1, otherwise. The nonnegative value 𝜆 is selected using the stability-based StARS^133^ method, using 200 subsamples, subsample size of 126 pseudobulk samples (63% of samples) and an instability threshold of 0.05. The adaptive LASSO strategy has several desirable properties: (1) Among many TFs with correlated TFAs, TFs with TFBS supporting a TF-gene interaction are prioritized (term 2). (2) Prior interactions not supported by the gene expression model are pruned (by term 1). (3) Prior information is not limiting, as “novel” TF-gene interactions, strongly supported by the gene expression model (term 1) are learned. This last property is important, given, for example, our incomplete knowledge of TFBS from ATAC-seq data.

##### Final GRN (model selection) and TFA estimates

To determine a confidence cutoff for the GRN, we evaluated out-of-sample gene expression prediction^59^. The confidence cutoff corresponds to model complexity (i.e., size or number of TF-gene regulatory interactions in the GRN) and is reported here as the average number of TF regulators per gene (**Fig. S2C**). For predictive performance evaluation, our gene expression samples were split into independent sets of training and test samples. Test performance was evaluated for three train-test splits of the data, where leave-out test data were: (1) all basal cell pseudobulk samples, (2) all samples from donor C or (3) ciliated pseudobulk samples at steady-state. Test performance was evaluated using R^2^-of-prediction (**R^2^_pred_**)), where 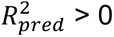 indicates that the GRN model improves prediction relative to the null model (in which, for each gene, the predicted gene expression is simply the mean observed across the training data samples). Across leave-out performance simulations, predictive performance (median **R^2^_pred_**) plateaued at 10 TFs/gene, justifying the selection of this model-size cutoff. These results were consistent for models based on TFA estimation using TF mRNA or prior-based TFA (**Fig. S2C**).

Thus, the two GRNs, resulting from models constructed using either TF mRNA or prior-based TFA and the full dataset, were integrated into an ensemble model (max combine of confidence ranks)^59^. To achieve a model size corresponding to an average of 10 TF per gene (as determined above), the highest-confidence 123,993 interactions were retained and re-ranked. The partial correlation between gene expression and TFAs were used to determine the sign of the interaction.

The TFA values reported were estimated using the final GRN (instead of the noisy prior) in **Equation 2**.

#### Gene regulatory network analysis

##### Differential TFA analysis

As TFA is a sum of gene expression values (**Equation 2**), we used two-tailed, paired t-tests (FDR = 5%) to identify TFs with differential TFA between two conditions. For TFs with dynamic TFA, we compared 2 or 6h to 0h, within a cell type. Analysis of basal, secretory and ciliated cells yielded 464 TFs with dynamic TFA in at least one cell type (**Table S3K**).

##### Definition of “core” TF regulators of steady-state cell type identity, using the GRN and HAE gene signatures

For each HAE cell type, “core” TFs were identified based cell type-associated TFA and their predicted regulation of signature genes in each cell type^59^ (**Fig. 1D, 1J**). Core TFs were classified as activators, if their predicted positive regulatory targets are enriched among the up-regulated signature genes, or, as repressors, if their repressed targets were enriched among the down-regulated signature genes (Fisher’s exact test, FDR = 5%). Putative activator and/or repressor TFs were subsequently filtered based on concordant, above-average TFA in the given cell type, relative to all HAE cell types. A subset of core TFs are highlighted (**Fig. 2A**, **2F**, **3A**, **3F**), while full results are available in **Table S3C, 3E-J**.

##### Definition of dynamic and baseline TF regulators of the IRG response

To distinguish TFs whose dynamic TFA drives IRGs from those that establish steady-state IRG expression, we used GSEA^134^ to test for the enrichment of a TF’s positive or negative target genes in the top or bottom of gene lists, ranked by log_2_FC gene expression, between 2 or 6h and steady state (0h), in each of three cell types: basal, secretory, ciliated (see **Enrichment Analysis** section below for technical details). Next, we enforced concordance between the TF’s regulatory effect and IFN-induced gene expression, retaining TFs with activating targets enriched among IIGs (normalized enrichment score (**NES**) > 0) or repressive targets enriched in IDGs (NES < 0). Our initial analysis was dominated by enrichment of canonical STAT1, STAT2 and IRF targets across all lineages, with few cell type-associated dynamic regulators identified. To uncover cell type-associated regulators of the cell type-associated IRGs, we repeated the analysis with genes from the “shared IIGs” and “shared IDGs” clusters in **Fig. 4A** excluded; these results are reported in panels i of **Fig. 6B, 6D** and **Table S5D**. We curated the list of 39 dynamic TFs shown in **Fig. 6B**: First we considered the top 40 activating and repressing TFs in each cell type (basal, ciliated and secretory) at either timepoint (2h and 6h), yielding a list of 108 TFs. Then we prioritized TFs whose enrichments were supported by concordantly dynamic, differential, TFA in the corresponding cell type(s). Some TFs were included in our final list of 39 despite not reaching statistical significance (ATF5, VEZF1 and ZNF467), which were included due to strong enrichment of their targets in IRGs along with a complementary trend in TFA (VEZF1 in ciliated, ZNF467 in secretory, and ATF5 in basal and secretory cells).

We also identified “baseline” TFs that set differential steady-state expression across cell types, also contributing to cell type-associated IRGs. Thus, we performed a second enrichment analysis, using Fisher’s exact tests (FDR=5%) to identify TFs with target enrichment in each IRG cluster (from **Fig. 4A**), separately testing TFs’ activating and repressed targets in the clusters’ IIGs and IDGs, respectively (panel iii, **Fig. 6B**, **6D**). (Note: although some IRG clusters are dominated by IIGs (shared IIGs, basal IIGs, ciliated-,secretory-associated IIGs) or IDGs (shared IDGs, basal IDGs, ciliated-, deuterosomal-associated, deuterosomal-associated and secretory-associated), each cluster contains a mix of IIGs and IDGs.) Only TFs with at least 5 predicted targets in the IRG cluster were considered in the analysis (**Table S5E**). Second, we required that candidate “baseline” TFs be supported by enrichment of their TFBS in the promoters of cell type-associated IRG cluster. We calculated the enrichment of TF motifs or *maxATAC*-predicted TFBSs ±2kb of IRG TSSs for each cluster (Fisher’s exact test, FDR = 5%, asterisks in panels iii, **Fig. 6B**, **6D**). TSSs were obtained from ENSEMBL v99 (January 2020). For genes with multiple known TSS, each TSS was treated independently, and a peak was associated with a gene if it was proximal to any gene TSS. We used *bedtools* v2.30.0^135^ *overlap* to find peaks within the gene TSS regions. Third, we required that baseline regulators be “core” TFs, as defined above.

TFs were manually clustered based on their TFA (panel ii, **Fig. 6B**, **6D**) and cell type-associated IRG enrichments (panel iii).

##### TF-TF module analysis

We calculated the overlap in predicted TF targets for each TF-TF pair, requiring concordance between the sign of regulatory interactions between pairs (e.g., both TFs positively regulate a gene would count as overlap, while discordant regulatory interactions (one TF positively but the other TF negatively regulates a gene) were not counted in the overlap. The analysis considered any TF with at least 20 gene regulatory interactions in which the absolute value of TF-gene partial correlation exceeded 0.01. In contrast the original implementation^59^, (1) both positive and negative regulatory interactions were considered (*calc_zscoredTfTfOverlaps* with *edgeOpt = ‘comb’*) and (2) the background-normalized score 𝑧_*ij*_, describing target overlap between TF*_i_* and TF*_j_*, newly considers mutually low overlap in addition to high overlap:

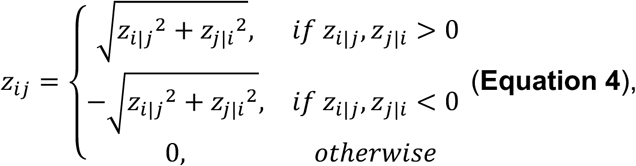

where 𝑧*_i_*_|*j*_ is the number of sign-concordant shared TF targets (overlaps) between TF*_i_* and TF*_j_*, z-scored according to the mean and standard deviation associated with the overlaps of TF*_j_* with the other TFs considered. The normalized overlap matrix was then filtered to include TFs with at least one significant overlap with another TF’s targets (FDR=5%). Then, we converted the filtered normalized overlap matrix to a distance matrix for hierarchical clustering using Ward distance. To select cluster number, we evaluated mean silhouette score, which plateaued, within error bars, at 90 clusters. We report the top-15 ranked^59^ TF-TF modules (**Table S4A**), with a subset visualized in **Fig. 2B**, **2H**, **3B**.

To implicate the TF-TF modules in coregulation of gene pathways, we defined the shared gene targets of a TF-TF module as those regulated by at least two TFs and used a Fisher’s exact test to assess overlap with GO biological processes. GO processes with at least 5 nominally expressed genes were included, and enrichments achieving P_adj_ < 0.05 are reported (**Table S4B**, **Fig. 2B**, **2H**, **3B**).

##### TF functional annotation predictions using the GRN and GO gene sets

Considering a TF’s predicted activating and repressive regulatory interactions separately, we functionally annotated TFs based on their predicted positive or negative regulation of GO biological processes. We limited our analysis to TFs with at least 5 positive or 5 negative interactions that were nominally expressed in the contexts considered (e.g., steady-state HAE cell type considered in **Fig. 2-3**). As above, GO processes with at least 5 nominally expressed genes were included. Enrichments achieving P_adj_ < 0.05 (Fisher’s exact test) are reported in (**Table S3D**).

We similarly annotated TF regulators of IFN-responsive GO biological processes (**Fig. S5**). For this analysis, we further limited the queried GO terms to include those enriched in any IRG cluster (**Fig. 4J**, **Table S3L**).

#### Generation of GRN subnetworks for visualization

##### Steady-state ciliated and deuterosomal GRN subnetworks

The selection of TFs and target genes (**Fig. 2C-E**) was focused on gene pathways associated with ciliated and deuterosomal cells. To identify pathways distinguishing deuterosomal and ciliated cells (**Fig. 2C, 2E**), we performed GSEA using GO biological processes on log_2_FC gene expression of deuterosomal with respect to ciliated cells at steady state and selected the top enriched pathways (NES > 0 for deuterosomal and NES < 0 for ciliated) (**Table S3M**). Leading edge genes were intersected with the respective cell-type gene signatures and the targets of the core TFs of each cell type (**Fig. 2A**). For the shared network (**Fig. 2D**), we performed GSEA for each cell type, using the mean of pairwise log_2_FC expression between ciliated or deuterosomal cells and each of the other HAE cell types. The top-4 shared, enriched pathways were selected, and leading edge genes were intersected with (1) shared signature genes of both cell types and (2) the targets of shared core TFs.

##### Steady-state ionocyte GRN subnetwork

To construct the ionocyte subnetwork (**Fig. 2G**), we performed GSEA using GO biological pathways on the average of pairwise log_2_FC gene expression between ionocytes and each of the other HAE cell types at steady state (FDR = 5%) (**Table S3N**). Top-enriched pathways, upregulated in ionocytes (“regulation of monoatomic ion transport”, “regulation of monoatomic ion transmembrane transport” and “monoatomic anion homeostasis”) were selected, and their leading-edge genes were intersected with ionocyte signature genes and the targets of ionocyte core TFs (**Fig. 2F**).

##### Steady-state basal and secretory GRN subnetworks

To construct the basal and suprabasal subnetworks at steady state (**Fig. 3C-D**), we identified basal-, suprabasal-unique, and shared gene signatures predicted to be regulated by at least two core TFs in the respective cell type(s). We ranked these targets based on the log_2_FC gene expression between basal and suprabasal cells and limited the networks to the top 30 differential targets. For the shared network (**Fig. 3E**), we ranked basal-suprabasal shared signature genes by calculating the mean of pairwise log_2_FC expression between basal or suprabasal cells and each of the other HAE cell types. The top 30 shared signature genes connected to shared core TFs (**Fig. 3A**) were selected

##### Steady-state secretory GRN subnetwork

We selected the top-8 secretory core TFs (panel i, **Fig. 3F**) and their interactions with secretory signature genes. To pare down the secretory signature genes, we selected the top 30, based on the average of pairwise log_2_FC gene expression between secretory cells and each other HAE cell type at steady state.

##### Subnetworks describing IRG regulation

For the regulation of shared IIGs (**Fig. 6C**), shared dynamic TFs (cluster B1, **Fig. 6B**) as well as select cell type-associated dynamic TFs, with strong enrichment (TFBS and target genes) in the shared IIGs (column “Shared IIGs” of panel iii, **Fig. 6B**), were included. For target gene selection, the union of top-10 highest confidence targets per TF were included. Due to the high degree of IRF7 and ISGF3 (i.e., large number of targets), the regulatory targets of these TFs are highlighted with node borders to improve legibility in the dense subnetwork.

Subnetworks (**Fig. 6E-G**) describe the regulation of cell type-associated IRGs in basal, secretory and ciliated cells, respectively. TFs are grouped according to classifications from **Fig. 6B**, **6D** and connected to the relevant IRG clusters in each cell type, with edges corresponding to enrichment (-log_10_(P_adj_)) of activating (red) or repressed (blue) targets in the IRG cluster. Furthermore, for TFs depicted as repressors of alternative cell-type IRGs, we verified, in a secondary analysis, that their TFBS were enriched proximally to the IRG clusters in the relevant cell types (**Fig. S4D-F**). For example, in the basal cell network, we confirmed, in a second analysis, that TF motif enrichment was supported using ATAC-seq peaks derived from basal cells specifically (**Fig. S4D**).

##### Chemokine and mucin subnetworks

We curated subnetworks explaining mucin (**Fig. 7A**) and chemokine (**Fig. 7C**) expression across HAE cell types. We attempted to identify at least one TF setting baseline and/or regulating dynamics (as applicable), to explain gene expression in each cell type. We prioritized TFs that: (1) were highlighted in **Fig. 6B, D** (available for basal, secretory or ciliated cells), (2) had high degree (number of mucin or chemokine targets), (3) congruence between TFA dynamics, sign of regulatory interaction and target gene expression.

#### Enrichment analyses

A combination of statistical approaches were used for enrichment analyses: (1) Fisher’s exact test for overrepresentation among sets (e.g., gene or peak clusters), (2) parameter-free gene set enrichment analysis (**GSEA**)^134^ for continuous results (e.g. *DESeq2*’s *ashr^125^*–shrunken log_2_FC estimates, as inputs to *fgsea* v1.24.0^136^, with the following options: ‘*gseaParam = 1*’, ‘*minSize = 5’*) and, in one context, (3) a simulation-based approach to control for chromatin accessibility in foreground and background peaks used for overrepresentation analysis. For all enrichment analyses, P_adj_ < 0.05 was used as a threshold for statistical significance.

##### Simulation-based TFBS enrichment analysis

To elucidate dynamic TF regulators of IFN-responsive peak clusters in each cell type (**Fig. 5A**), we designed a simulation-based TFBS enrichment analysis to identify TFs with over- or under-representation of TFBS in IFN-responsive peaks, while controlling for baseline accessibility in the given cell type. (Controlling for baseline accessibility was important, as most cell type-associated IFN-responsive peak clusters had elevated baseline accessibility in the corresponding cell types (clusters 2-5, **Fig. 5A**, **S3D**, **S3H**) and we wanted to distinguish TFs that drive cell type-specific chromatin opening from those that establish the baseline chromatin landscape.) For each cell type, we compiled peaks detected at any time point in the cell type and split them into IFN-responsive (overlapping peaks in **Fig. 5A**) and non-dynamic (“background”) sets. Then, for each IFN-responsive peak (“IRP”) cluster (from **Fig. 5A**), we identified TFBS occurrences in our cluster of interest (“IRP” set) and background peaks, using *maxATAC* predictions (if available) and motif scanning otherwise (see above), determining the number of base pairs (**bp**) overlapping a predicted TFBS. For each IRP cluster, in a given cell type, the odds ratio (**OR**) was calculated, using the IRP cluster as the foreground set:

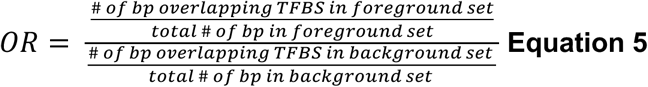

To determine significance: For each IRP cluster (from **Fig. 5A**) in each cell type, we generated a “target” distribution of donor-averaged steady-state VST counts, and these counts were binned into deciles. We then generated 200 random samples without replacement from the background peaks, where each random sample was matched to the IRP set in terms of: number of peaks and the VST count distribution (matching deciles). TFBS were identified in our random peak samples, and “random 𝑂𝑅_*i*_” was calculated for random sample *i* substituting the random sample as the foreground set in **Equation 5**.

From the 200 random samples, an empirical null distribution of ORs was determined for each TF, IRP cluster and cell type. We fit the null distribution assuming a normal distribution, using the inverse normal CDF to estimate two-sided p-values for the observed OR. This analysis was performed using the *GenomicRanges* v1.56.2^137^ package (**Fig. 5H-M, Table S6C**).

##### Enrichment of gene and peak sets (Fisher exact test or GSEA)

For enrichment analyses involving gene *and* peak features, peaks were associated with genes based on proximity (± 2kb) to the gene’s TSS. TSSs were obtained from ENSEMBL v99 (January 2020). For genes with multiple known TSS, each TSS was treated independently, and a peak was associated with a gene if it was proximal to any gene TSS. Similarly, peaks that were located proximal to multiple genes were independently associated with each gene. We used *bedtools* v2.30.0^135^ *overlap* to find peaks within the gene TSS regions.

Here we summarize sets used for enrichment analyses (if not already described above in the “GRN Analysis” section):

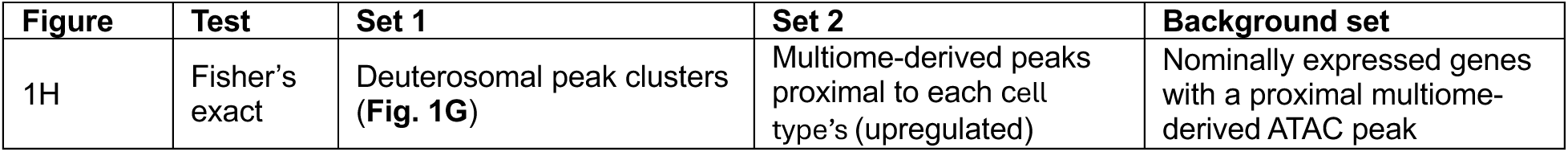
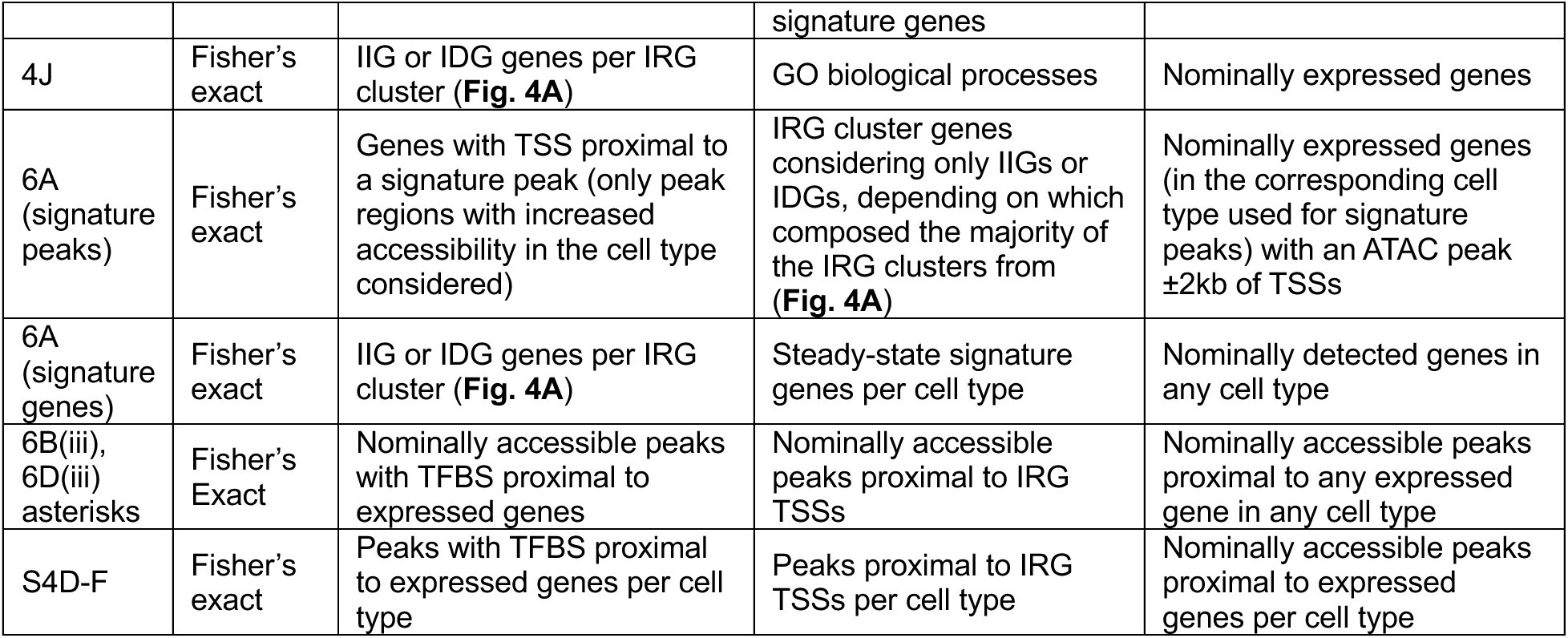

Here we summarize sets used for enrichment analyses involving the GRN (with more details regarding rationale in the GRN Analysis section above):

| Figure | Test | Set 1 | Set 2 | Background set |
| --- | --- | --- | --- | --- |
| 2A, 2F, 3A, 3F | Fisher's exact | TF activating or repressing targets from the GRN | Cell type signature genes | Cell type-specific nominally expressed genes |
| 2B, 2H, 3B | Fisher's exact | Shared TF targets, predicted to be regulated by at least two TFs in module | GO biological processes | Nominally expressed genes in any cell type |
| 2C-E, 2G, 3C-E, 3G | GSEA | log2FC (cell type differences in gene expression at steady state) | GO biological processes | Nominally expressed genes in any cell type |
| 6B (i), 6D (i) | GSEA | log2FC of 2 or 6h v. 0h expression, in a given cell type | TF activating or repressing targets from the GRN | Nominally expressed genes in any cell type |
| 6B(iii), 6D(iii), 6C, 6E-G | Fisher's exact | TF activating or repressing targets | IIG or IDG genes per IRG cluster (**Fig. 4A**) | Nominally expressed genes in any cell type |
| S5 | Fisher's exact | TF activating or repressing targets | GO biological processes enriched in IRG clusters (**Fig. 4J**) | Nominally expressed genes in any cell type |

#### Single cell analyses

##### Cell cycle scoring

Cell cycle scoring was performed using the *CellCycleScoring* function provided in *Seurat* v4.3.0^109^, assigning the phase with the highest score to each cell (**Fig. S1E**).

##### Per-cell peak activities

To examine the relative single-cell accessibility of interferon-responsive peaks, we used *chromVar*^138^ to calculate the per-cell accessibility (deviation) score for each peak relative to the background reference set of peaks. The resulting peak x cell deviation matrix was z-scored and barcodes corresponding to each cell type and timepoint were aggregated for visualization (**Fig. S3D**).

#### Clustering and data visualization

The k-means algorithm was used for clustering (**Fig. 1C-D, 1G, 1J, 4A-B, 5A**). Heatmaps were visualized with *ComplexHeatmap* v2.14.0^139^ or *ggplot2* v4.0.1^140,141^ package. All GRNs were visualized using the desktop version of *Cytoscape* v3.10.4^142^.

### ADDITIONAL RESOURCES

The resources developed in our study have been made interactively available from https://github.com/MiraldiLab/airwayGRN. These resources include: (1) *GRN Visualization:* All subnetworks described can be interacted with through *Cytoscape* v3.10.4 by loading Cytoscape session files. (2) *Genome browser of accessible chromatin and maxATAC TF binding site predictions*: A track data hub^144^ was generated to visualize both chromatin accessibility data and the maxATAC predictions in the UCSC Genome Browser (see the “*Prediction of TFBS using maxATAC neural network models*” section). Predictions for all *maxATAC* TF models were aggregated per cell type-timepoint (e.g., basal cells at 0h) into BED files across all TFs, resulting in a total of 9 BED files. These BED files were then converted into bigBED format using the *bedToBigBed* utility from *UCSCtools* v466^145^, and the data hub (https://gb.research.cchmc.org/hub/group/HAE/hub.txt) was compiled following the format described in the Basic Track Hub Quick Start Guide: (https://genome.ucsc.edu/goldenPath/help/hubQuickStart.html). (3) *Codebases*: The codebases to generate all figures in the manuscript are freely available from the repository as well.

---

**Table S1.** Pseudobulk gene expression and steady-state signature genes. Batch-corrected, normalized pseudobulk gene expression matrices from the scRNA-seq dataset for (**A**) data visualization and (**B**) GRN inference. Signature genes at steady-state for the major HAE cell types are annotated in (**A**) (*DESeq2*, |log_2_FC| > 0.58, P_adj_ < 0.1). Related to Figures 1, 3, 7 and S4.

**Table S2.** Pseudobulk chromatin accessibility and signature peaks. Batch-corrected, normalized pseudobulk chromatin accessibility matrices from the (**A**) snATAC-seq and (**B**) sn-Multiome-seq datasets. Signature peaks at steady-state for the major HAE cell types (*DESeq2*, |log_2_FC| > 0.58, P_adj_ < 0.1) and genes proximal to peaks (±2Kb of TSS or ±10Kb of gene bodies) are annotated in (**A**). Related to Figures 1, 5, S1 and S3.

**Table S3.** Gene regulatory network and TF target enrichment. (**A**) GRN-predicted TF-target interactions with reported stability scores and correlation of target gene expression. (**B**) Transcription factor activity (TFA) estimates. Enrichment of TF activating or repressing interactions in (**C**) steady-state gene signatures and (**D**) GO biological processes (Fisher’s exact test). (**E-J**) Steady-state core networks for basal, suprabasal, ciliated, deuterosomal, ionocytes and secretory cells. (**K**) Significance of the change in TFA (2 or 6h relative to 0h, based on t-test). (**L**) Enrichment of TF activating or repressing targets in IFN-responsive GO biological processes (Fisher’s exact test). GSEA of GO biological processes in the differential expression profiles of (**M**) deuterosomal and ciliated cells or (**N**) ionocytes relative to the major HAE cell types. Related to Figures 1-3, 5-7, S2 and S4-5.

**Table S4.** TF-TF module analysis and enrichment of shared targets in GO biological pathways. (**A**) Top 15 TF-TF modules for steady-state TFs. (**B**) Enrichment of shared TF targets in GO pathways (Fisher’s exact test, P_adj_ < 0.05). Related to Figures 2 and 3.

**Table S5.** Cell type-specific interferon responsive genes (IRGs) and functional enrichments. (**A**) IRG identification using differential gene expression for each major HAE cell type at 2 or 6h relative to steady state (*edgeR*). (**B**) IRG cluster assignments based on cell type-and timepoint-associated expression dynamics. Enrichment of (**C**) GO biological processes in IFN-increased or decreased IRG subclusters and (**D**) TF activating or repressing targets in IRG subclusters, respectively (Fisher’s exact test). (**E**) GSEA of TF activating or repressing targets in cell type-specific differential gene expression profiles (log_2_FC at 2 or 6h relative to steady-state). IRGs belonging to the shared IIG and IDG clusters were omitted. Enrichment of IRG subclusters in (**F**) steady-state signature genes or (**G**) genes proximal (±2Kb of TSS) to steady-state signature peaks (Fisher’s exact test). Related to Figures 4-7 and S3-4.

**Table S6.** Cell type-specific interferon responsive peaks (IRPs) and TFBS enrichments. (**A**) IRP identification using differential analysis of chromatin accessibility for each major HAE cell type at 2 or 6h relative to steady state (*edgeR*, P_adj_ < 0.05 and |log_2_FC| > 1). (**B**) IRP cluster assignments based on cell type-associated accessibility dynamics. Enrichment of *maxATAC*-predicted TF binding sites or motif occurrences in (**C**) IFN-increased peak clusters, controlling for cell type-specific steady-state accessibility (simulation-based empirical p-value estimation), and in (**D**) reference peaks or (**E**) cell type-specific peaks proximal (±2Kb of TSS) to IRG subcluster genes (Fisher’s exact test). Related to Figure 5-6 and S4.

## Notes

### Competing Interest Statement

BRR is on the Scientific Advisory Board for Dispatch Biotherapeutics.

